# High-Throughput FRET Affinity Screening Technique (HTFAST) For Cell-Free Expressed Binding Protein Characterization

**DOI:** 10.64898/2026.02.12.697512

**Authors:** Sepehr Hejazi, Kimia Noroozi, Vito Jurasic, Laura R Jarboe, Nigel F Reuel

## Abstract

The rapid engineering of high-affinity binding proteins, such as nanobodies and single-domain antibodies (sdAbs), is increasingly driven by cell-free, machine–learning–guided optimization. However, high-throughput, quantitative characterization of binding affinity remains a major bottleneck, particularly for proteins expressed in cell-free systems without purification. Here, we present High-Throughput FRET Affinity Screening Technique (HTFAST) for rapid affinity characterization of binders expressed directly in crude *E. coli* cell-free protein synthesis reactions. HTFAST leverages Förster resonance energy transfer (FRET) between fluorescent-protein–fused binders and dye-labeled antigens to enable real-time, quantitative measurement of equilibrium dissociation constants. We systematically optimized fluorophore pairs used and labeling parameters using the SpyTag003–SpyCatcher003 model system. Using donor-quenching and acceptor-emission FRET analyses, HTFAST reliably quantified nanomolar binding affinities in crude lysates for SpyTag003–SpyCatcher003 model system. We validated the platform for nanobodies by characterizing a CD4-binding nanobody, Nb457, and benchmarking multiple SARS-CoV-2 receptor-binding domain sdAbs, demonstrating HTFAST’s ability to rank binding strengths across a range of affinities. Finally, we demonstrate that both binding partners can be expressed directly in CFPS, further streamlining screening workflows. Overall, HTFAST provides a scalable, quantitative, and cell-free–compatible approach for high-throughput affinity screening, well suited for DBTL campaigns aimed at accelerating the development of next-generation binding proteins.

## Introduction

Antibodies and their variants have dominated protein research and commercialization due to their pivotal role in health and their applications in therapeutics and diagnostics.^1^Small affinity proteins, including antibody fragments and nanobodies (single variable domains of heavy-chain antibodies, VHHs), have emerged as promising alternatives to traditional full scale antibodies for accelerated design–build–test–learn (DBTL) strategies in protein engineering.^2–4^These compact proteins (∼15 kDa) combine high affinity for target antigens with enhanced stability, improved tissue penetration, and facile expression across diverse expression systems.^5,6^

Traditional nanobody discovery through in vivo immunization of camelids is time-consuming, costly, and limited by the direction of host immune response. In vitro selection platforms such as phage, yeast, and ribosome display offer attractive alternatives by enabling discovery from synthetic libraries and isolating high-affinity binders without the need for animal immunization.^7,8^Although these display-based methods are powerful for initial discovery, they are less compatible with AI-driven protein engineering workflows because they sample an enormous and largely unconstrained sequence space, providing limited control over the specific variants generated and evaluated.^9,10^Moreover, the data these screens return is limited to the high-affinity variants, and little is learned from other sequences.

To address this limitation, cell-free, machine-learning–guided platforms have emerged as rapid protein synthesis systems that support accelerated DBTL cycles.^11^By leveraging only the transcriptional and translational machinery extracted from cells, cell-free protein synthesis (CFPS) enables the production of functional proteins directly from synthetic gene templates in under 24 h, making it well-suited for iterative DBTL workflows.^12,13^CFPS has been demonstrated as a powerful platform for expressing not only nanobodies but also full-length monoclonal antibodies.^14–16^However, the major bottleneck in high-throughput, CFPS DBTL campaigns remains the screening of thousands of expressed nanobody variants and the acquisition of high-quality, quantitative data suitable for machine-learning workflows.^17^

A brief review of current affinity characterization methods (Table 1) will help frame the utility of the approach presented in this paper. First, ELISA assays have been widely used in plate-based formats. However, ELISA provides limited quantitative information and relies on multiple washing steps, a restricted dynamic range, and serial dilutions, all of which constrain its throughput.^18^Techniques such as surface plasmon resonance (SPR) and biolayer interferometry (BLI) are considered gold standards for obtaining binding kinetic data,^19,20^but their throughput is inherently limited, and they are not compatible with large-scale plate-format screening. Furthermore, the complex background of CFPS, particularly in crude lysate systems, necessitates additional protein purification steps prior to characterization, further reducing experimental throughput.^21^

**Table 1.**
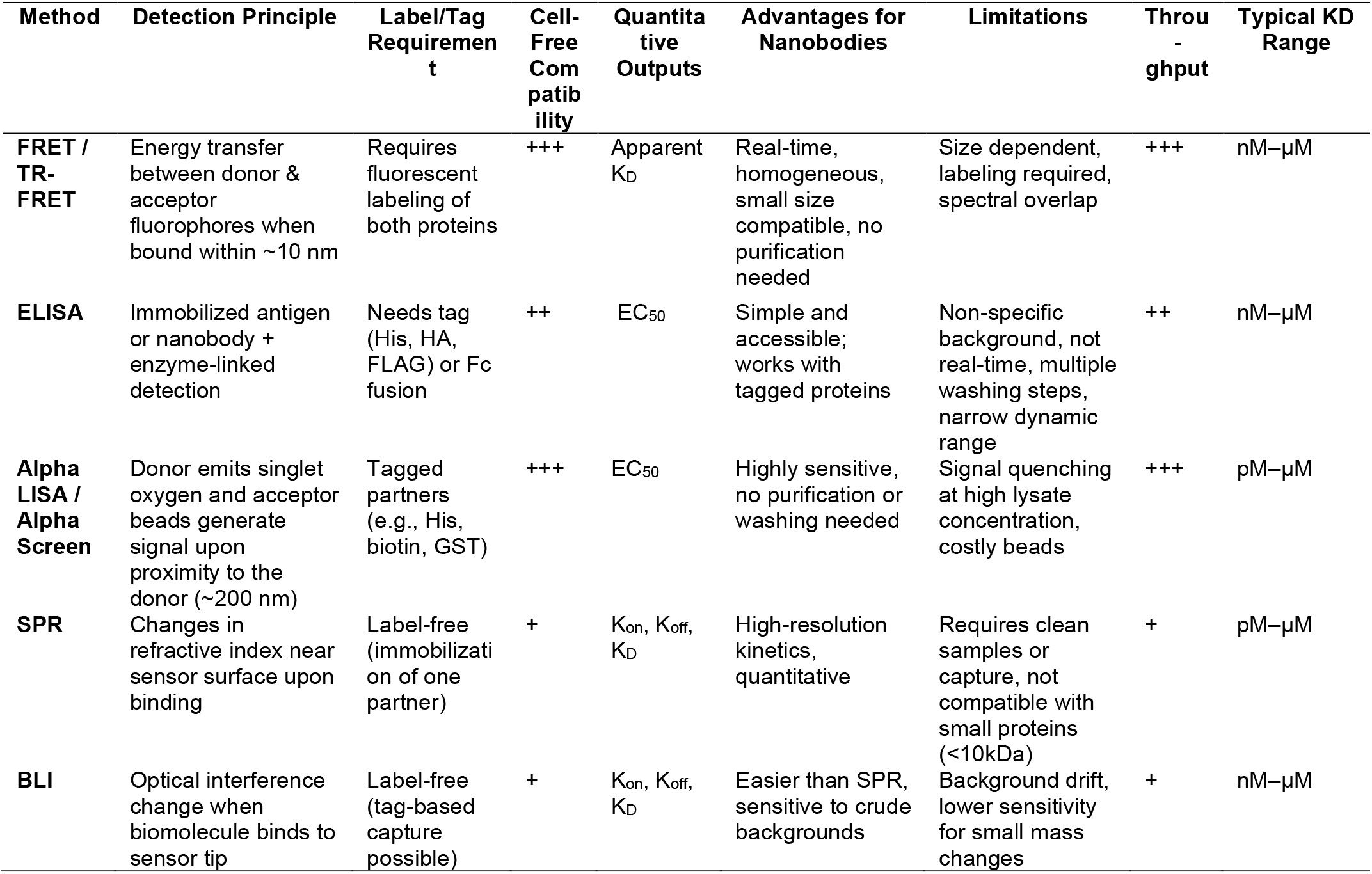
Comparison of current methods for affinity characterization and screening

| Method | Detection Principle | Label/Tag Requirement | Cell-Free Compatibility | Quantitative Outputs | Advantages for Nanobodies | Limitations | Throughput | Typical KD Range |
| --- | --- | --- | --- | --- | --- | --- | --- | --- |
| **FRET / TR-FRET** | Energy transfer between donor & acceptor fluorophores when bound within ~10 nm | Requires fluorescent labeling of both proteins | +++ | Apparent $K_D$ | Real-time, homogeneous, small size compatible, no purification needed | Size dependent, labeling required, spectral overlap | +++ | nM–μM |
| **ELISA** | Immobilized antigen or nanobody + enzyme-linked detection | Needs tag (His, HA, FLAG) or Fc fusion | ++ | $EC_{50}$ | Simple and accessible; works with tagged proteins | Non-specific background, not real-time, multiple washing steps, narrow dynamic range | ++ | nM–μM |
| **Alpha LISA / Alpha Screen** | Donor emits singlet oxygen and acceptor beads generate signal upon proximity to the donor (~200 nm) | Tagged partners (e.g., His, biotin, GST) | +++ | $EC_{50}$ | Highly sensitive, no purification or washing needed | Signal quenching at high lysate concentration, costly beads | +++ | pM–μM |
| **SPR** | Changes in refractive index near sensor surface upon binding | Label-free (immobilization of one partner) | + | $K_{on}$, $K_{off}$, $K_D$ | High-resolution kinetics, quantitative | Requires clean samples or capture, not compatible with small proteins (<10kDa) | + | pM–μM |
| **BLI** | Optical interference change when biomolecule binds to sensor tip | Label-free (tag-based capture possible) | + | $K_{on}$, $K_{off}$, $K_D$ | Easier than SPR, sensitive to crude backgrounds | Background drift, lower sensitivity for small mass changes | + | nM–μM |

Next, AlphaLISA and AlphaScreen are promising proximity-based assays for characterizing protein–protein interactions (PPIs) in unpurified cell-free expression systems and have been successfully applied in high-throughput screening (HTS) formats.^21,22^However, their broader adoption is constrained by the high cost of specialized reagents, special plate-reader specifications, and their limitation to endpoint measurements that do not provide kinetic PPI information.^23^

Förster resonance energy transfer (FRET) based assays have been widely used to obtain kinetic information on protein–protein interactions (PPIs) in complex biological environments. FRET relies on the non-radiative transfer of energy from an excited donor fluorophore to a nearby acceptor fluorophore when the two are within close proximity, typically less than 10 nm. ^24,25^This makes it an ideal screening platform for monitoring PPI between small binding proteins such as nanobodies and purification tags.

Here, we describe a High-Throughput FRET Affinity Screening Technique (HTFAST) for rapid quantification of kinetic parameters of binding proteins expressed in *E. coli* cell-free systems. HTFAST utilizes FRET between a fluorescent-protein–labeled fusion protein and its binder interacting with a labeled antigen, enabling real-time, homogeneous, high-throughput characterization of binding affinity and kinetics (**Figure 1**).

**Figure 1.**
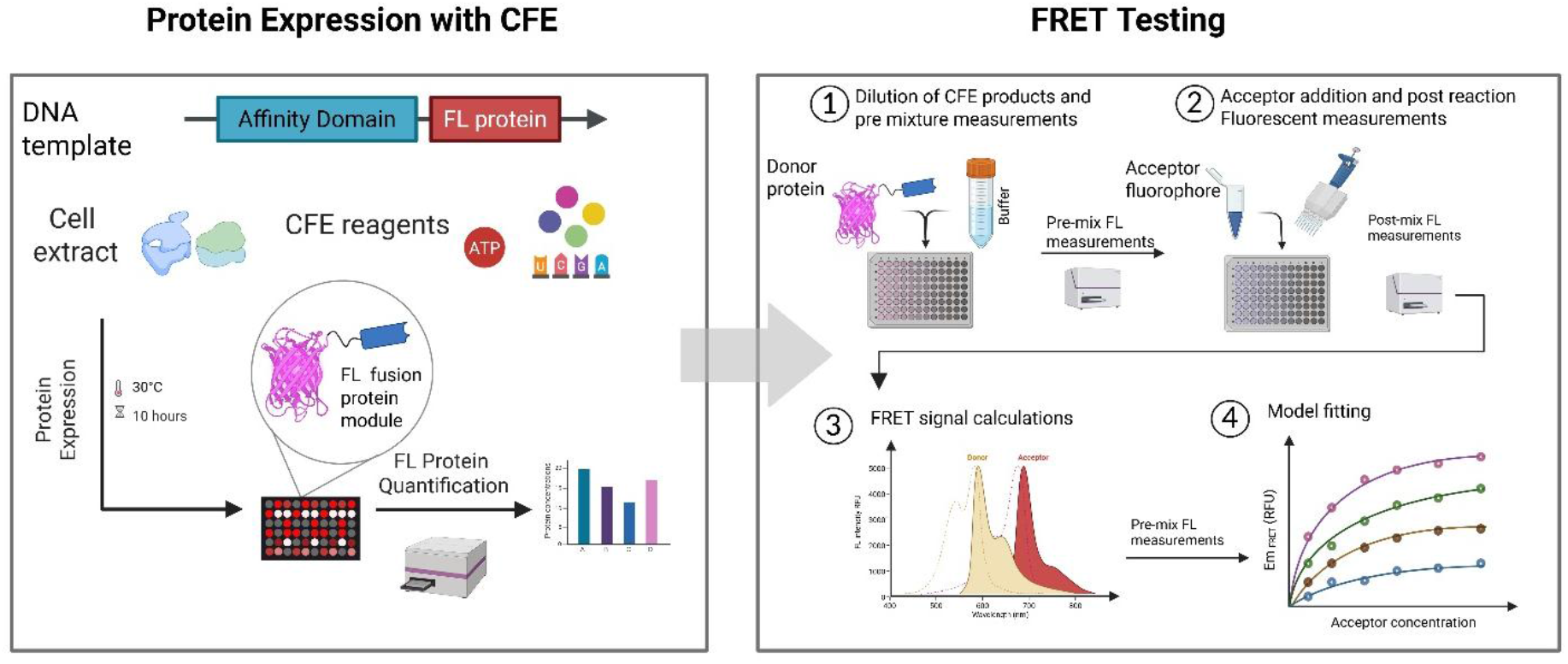
Workflow of cell-free protein affinity quantification by HTFAST. FL-protein fusions are expressed in crude lysates, and FRET-based kinetic measurements are performed by stepwise addition of acceptor-conjugated proteins. Fluorescence readouts before and after each addition are used to calculate FRET signals, which are analyzed to determine kinetic parameters.

## Materials and Methods

### Materials

SpyCatcher003 (Bio-Rad Laboratories), Alexa Fluor 647 NHS ester (Thermo Fisher Scientific), ATTO 532 NHS ester (Thermo Fisher Scientific), ATTO 594 NHS ester (Millipore Sigma), Zeba Spin Desalting Columns, 7 kDa MWCO (Thermo Fisher Scientific), Human CD4/LEU3 Protein, His-tag (Acro Biosystems), and SARS-CoV-2 Spike RBD Protein (Acro Biosystems) were used as received.

### Protein dye-labeling

Protein conjugation with NHS-ester fluorophores followed the manufacturer’s recommended procedures. Briefly, dyes were dissolved in anhydrous dimethyl sulfoxide (DMSO) at a concentration of 10 mg/mL. Unless otherwise specified, proteins were reacted with dye at a 5:1 molar ratio in 1 mL of 0.1 M sodium bicarbonate buffer (pH 8.3) and incubated for 1 h at room temperature. To generate samples with varying degrees of labeling (DOL), dye-to-protein molar ratios of 1:1, 2:1, 4:1, and 8:1 were also tested.

Reactions were quenched by adding 0.1 mL of freshly prepared 1.5 M hydroxylamine. Excess, unreacted dye was removed using 7 kDa Zeba spin desalting columns. Protein concentrations were determined using the Qubit Protein Assay.

The degree of labeling (DOL) was calculated from absorbance measurements using:

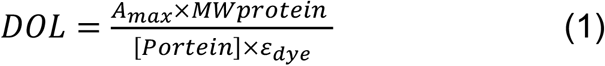

where A_max_is the absorbance of the dye at its peak wavelength, *MW*_protein_is the molecular weight of the protein, ε_dye_is the dye’s molar extinction coefficient at its absorbance maximum, and [Protein] is the protein concentration in mg/mL.

### Cell lysate preparation

Cell lysate for cell-free protein synthesis was generated using *E. coli* One Shot™ BL21 Star™ (DE3) (Thermo Fisher Scientific). Cultures were grown following established methods^26^and cell pellets were flash-frozen in liquid nitrogen, then stored at −80°C until use. For lysate preparation, frozen pellets were thawed on ice and resuspended in ice-cold S30 acetate buffer (10 mM Tris-acetate, pH 8.2; 14 mM magnesium acetate; 60 mM potassium acetate) at a ratio of 1 mL buffer per gram of wet cell mass. Cells were disrupted using an Avestin EmulsiFlex-B15 homogenizer operated at approximately 24,000–26,000 psi. The homogenate was clarified through two sequential centrifugation steps (12,000 ×g, 4°C, 10 min each) to remove cell debris. The final clarified lysate was aliquoted, rapidly frozen in liquid nitrogen, and stored at −80°C for downstream cell-free expression reactions.^27^

### Cell-free expression reactions

A previously validated energy mix formulation^28^was used to support transcription and translation in the CFE reactions. To ensure the proper formation of disulfide bonds, 4 mM oxidized glutathione (GSSG), 1 mM reduced glutathione (GSH) was added to the energy mixuture solution.^29^Each reaction was assembled by combining lysate (24% v/v) with the energy mix (33% v/v), along with 5 nM DNA template. Nuclease-free water was added to bring the total reaction volume to 15 μL per well. To ensure the proper formation of disulfide bonds, 4 mM oxidized glutathione (GSSG), 1 mM reduced glutathione (GSH) was added to the energy mixuture solution.

### Fluorescent Protein Concentration Measurements

To generate standard curves for quantifying fluorescent protein concentrations, purified mScarlet3 and mCitrine were obtained using a previously established Strep-tag II purification workflow.^30^Proteins were isolated with Strep-Tactin® XT 4Flow® High-Capacity Spin Columns (IBA Lifesciences) following the manufacturer’s instructions. Purified protein concentrations were determined using the Qubit Protein Assay Kit. Known amounts of each fluorescent protein were then added to cell-free reaction mixtures at defined dilution ratios to construct calibration curves (Supporting Figures S1,S2).

### FRET Assay Setup and Fluorescence Measurements

FRET measurements were carried out using a two-dimensional titration matrix of donor and acceptor modules. Three or four serially diluted concentrations of the donor protein were dispensed into a 384-well black, flat-bottom plate (Corning), followed by the addition of serially diluted acceptor protein to generate the full 2D concentration matrix. Fluorescence was recorded both before and after the addition of the acceptor. Three channels were measured: FL_DA_ (donor excitation, acceptor emission), FL_DD_ (donor excitation, donor emission), and FL_AA_ (acceptor excitation, acceptor emission). Absorbance at the relevant excitation and emission wavelengths was also collected to enable inner filter effect correction using the Lakowicz method.^31^

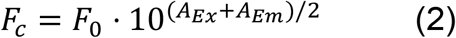

Where F_C_ is the corrected fluorescence, F_0_ is the raw fluorescence, A_Ex_ is the absorbance at the excitation wavelength and A_Em_ is the absorbance at the emission wavelength.

For the acceptor-emission method, the FRET-specific signal was calculated using:

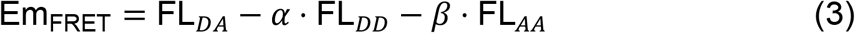

where **α** represents donor bleed-through, defined as the ratio of FL_DA_ to FL_DD_ measured from donor-only samples (prior to acceptor addition). **β** represents acceptor direct excitation, calculated as the ratio of FL_DA_ to FL_AA_ from acceptor-only samples (acceptor added to buffer-only wells).

For the donor-quenching approach, FL_DD_ was measured before and after addition of the acceptor. Background donor-channel fluorescence arising from the acceptor (FL_DD_ of acceptor-only wells) was subtracted prior to fitting.

### Model fitting and KD calculations

For analysis of acceptor-emission data, we applied a quadratic 1:1 binding model appropriate for conditions where neither binding partner is in large excess.^32^The FRET signal was modeled as (See Supporting Information Section 2 for how the equations are derived):

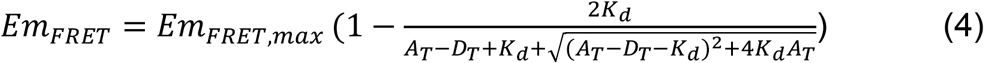

where Em_FRET_ represents the corrected acceptor-emission FRET intensity, and *Em*_FRET,max_ corresponds to the maximal achievable FRET signal when all donor molecules are bound to acceptor. In this expression, D_T_ is the total donor concentration in the assay, and A_T_ denotes the titrated acceptor concentration. A custom Python script (Supporting Information section 6) was developed to fit the acceptor-emission data to this quadratic binding model, using K_d_ and *Em*_FRET,max_ as free parameters to extract the equilibrium dissociation constant.

For the Donor quenching approach, we used the same quadratic binding model but for the donor quenching approach (See supporting information section 2 for how the equations is derived):

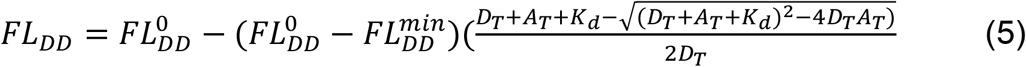

Where ΔFL_DD_is the Donor fluorescence after the acceptor is added, 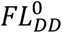 is donor signal with no acceptor, 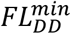 is donor signal at saturating acceptor. A custom Python script (Supporting file 2) was developed to fit the donor-quenching data to this quadratic binding model, using K_d_ and 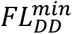 as free parameters to extract the equilibrium dissociation constant.

For Fret efficiency (E),donor fluorescence was recorded before and after saturation with the acceptor protein module^33^, and FRET efficiency was calculated as:

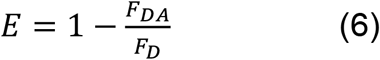

where F_D_ is the donor fluorescence in the absence of acceptor and F_DA_ is the donor fluorescence after acceptor binding.

## Results and Discussion

### Fluorophores selection and optimization using Spy-Catcher003 and cell-free expressed SpyTag003 system

The principle of HTFAST relies on the affinity interaction between a fluorescent-protein (FL)–labeled binder serving as the FRET donor and a labeled antigen serving as the acceptor. Because both interaction partners must carry fluorophores, we expressed the donor protein module as a fluorescent-protein fusion in a cell-free protein synthesis (CFPS) system. This strategy enables real-time quantification and monitoring of protein expression in crude lysate, which is essential for HTFAST measurements.^34^Also, because many antigens for these binders are commercially available, naturally extracted, or difficult to express in cell-free systems, we focused on dye conjugation of procured antigens. This approach enables FRET between protein-labeled CFPS-expressed binders and dye-conjugated antigens.

To evaluate different FL protein candidates, we first employed the SpyTag003– SpyCatcher003 system as the binding pair (**Figure 2a**). The short 16–amino acid SpyTag003 peptide binds tightly to its partner, SpyCatcher003, enabling efficient FRET at a short donor–acceptor distance. A previous study used an mClover–SpyTag003 donor with Alexa Fluor 555–labeled SpyCatcher in a stopped-flow assay to determine the association and dissociation rate constants for this interaction.^35^

**Figure 2.**
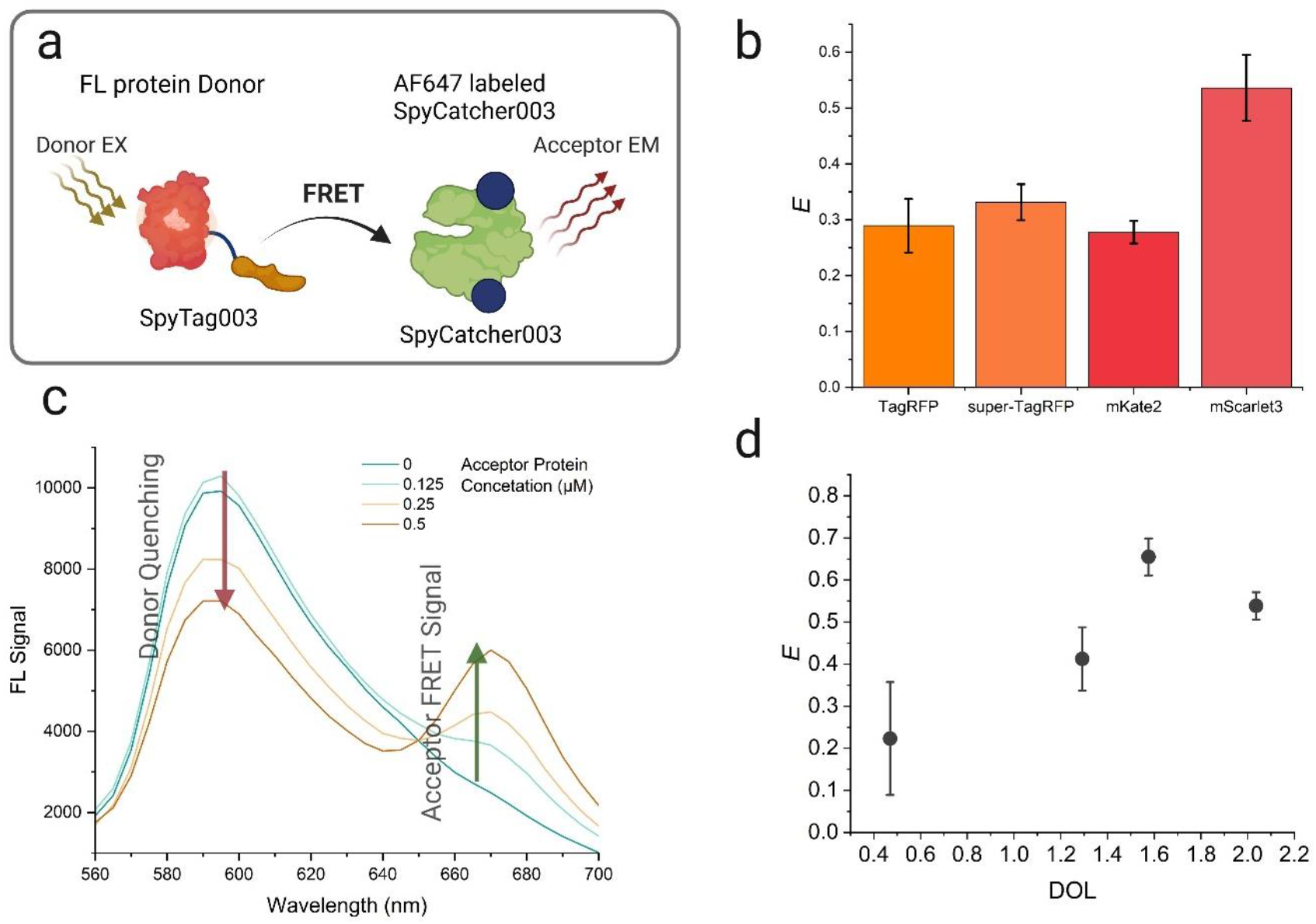
a) Schematic of the FRET interaction between the fluorescent protein–labeled SpyTag003 donor module and the Alexa Fluor 647–conjugated SpyCatcher003 acceptor module. b) Emission spectral changes of the mScarlet3–SpyTag003 donor upon different concentrations of AF647–SpyCatcher003. c) FRET efficiency of different fluorescent proteins used as donors with AF647–SpyCatcher003. Bars represent the mean of three independently prepared cell-free reactions; error bars indicate standard error. d) FRET efficiency of mScarlet3–SpyTag003 paired with AF647–SpyCatcher003 prepared at different DOL. Bars represent the mean of three independently prepared cell-free reactions; error bars indicate standard error

To label the SpyCatcher003 antigen, NHS-ester dyes provide a simple and rapid method to label primary amines (exposed lysine residues), which, although non-specific, offers an efficient way to conjugate dyes to proteins. Initially Atto 594 was tested as an acceptor paired with an mScarlet3 donor within the same system. However, this combination again failed to produce robust FRET signals, likely due to substantial spectral crosstalk caused by close excitation and emission overlap between the fluorophores.

The Alexa Fluor 647 (AF647) dye was selected to minimize interference from the intrinsic autofluorescence of E. coli–based cell-free expression systems.^36,37^To select suitable donor proteins, it is essential that the donor emission spectrum overlaps with the excitation spectrum of Alexa Fluor 647. This makes red and orange fluorescent proteins ideal candidates due to their spectral properties. We selected four fluorescent proteins, TagRFP (Ex λ = 555 nm, Em λ = 584 nm), SuperTagRFP (Ex λ = 555 nm, Em λ = 579 nm), mKate2 (Ex λ = 588 nm, Em λ = 633 nm), and mScarlet3 (Ex λ = 569 nm, Em λ = 592 nm),and evaluated their FRET efficiency (*E*) to Alexa Fluor 647 using donor quenching (**Figure 2b**).

We observed that mScarlet3 exhibited the best performance, achieving a maximum FRET efficiency of 53%, and therefore selected it as the donor fluorophore for subsequent experiments (**Figure 2c**). This is consistent with the higher quantum yield of mScarlet3 (0.75)^38^compared to mKate2 (0.4)^39^, Tag-RFP (0.48)^40^and super-Tag-RFP (0.53)^41^. It should be noted that CyOFP1 (Ex λ = 497 nm, Em λ = 589 nm) with quantum yield of 0.76 ^42^was also used as donor protein but no FRET signal was observed when paired with Alexa Fluor 647 conjugated acceptor likely due to its lower spectral overlap with AF647 (J(λ) = 11.83 × 10^15^ M^−1^ cm^−1^ nm^4^), compared with the substantially higher overlap for mScarlet3 (J(λ) = 15.24 × 10^15^ M^−1^ cm^−1^ nm^4^), as calculated using the FPbase FRET Calculator. ^43^

We have also evaluated another FRET strategy in which Atto 532–labeled SpyCatcher003 was tested as a donor with SpyTag003 fused to various fluorescent protein acceptors, but reliable FRET signals could not be obtained. Significant spectral bleed-through was observed, likely arising from residual excitation overlap (“left shoulder”) in the excitation spectra of fluorescent proteins.^38^Because of this, fluorescent proteins function more effectively as donors to minimize non-specific excitation and bleed-through.

To further optimize the FRET system, we examined the effect of the degree of labeling (DOL) of Alexa Fluor 647 on SpyCatcher003. Various dye-to-protein ratios were used during conjugation, and unreacted dye was removed by purification through desalting columns. DOL values were determined from the absorbance of the dye and the protein. We observed that FRET efficiency increased with higher DOL and reached a maximum at a DOL of 1.6 (E = 70%), after which additional dye conjugation led to a decline in performance (**Figure 2d**). This behavior can be explained by insufficient dye molecules at low DOLs, where some binding events fail to quench donor fluorescence, and by excessive labeling at high DOLs, which may sterically hinder or partially obstruct the antigen-binding interface, reducing effective binding affinity and consequently lowering FRET efficiency.

### Kinetic Analysis of Protein Binding Using FRET

To apply our FRET system for screening affinity proteins such as nanobodies and binding tags, we used a quantitative FRET approach to determine equilibrium dissociation constants (K_D_) for each interaction. Two quantitative approaches are commonly used for determining K_D_; acceptor-emission analysis (Figure 3a,b) and donor-quenching analysis (Figure 3c,d). In the acceptor-emission method, the FRET signal is measured as the increase in acceptor fluorescence upon excitation at the donor excitation wavelength. Titrating varying concentrations of donor protein with a series of acceptor concentrations generates equilibrium binding curves (Figure 3a). A global quadratic binding model is then fitted to all curves simultaneously to obtain a single global K_D_, minimizing the residual error across all acceptor concentrations. Lower K_D_ values produce steeper binding curves, reflecting tighter interactions (Figure 3b).

**Figure 3.**
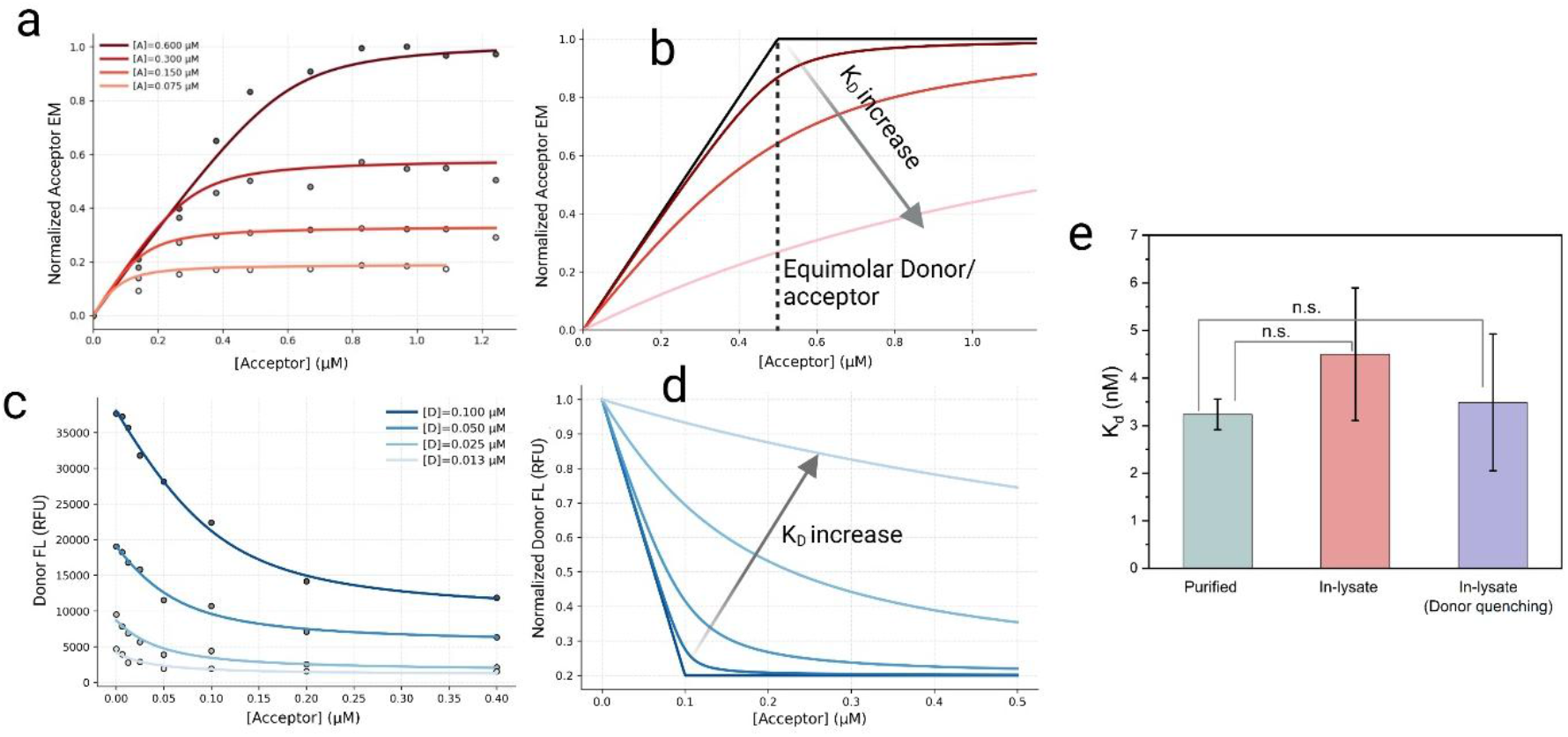
a) Acceptor emission (FRET) response of mScarlet3–SpyTag003 (donor, Protein A) titrated with AF647– SpyCatcher003 (acceptor, Protein B) at varying donor concentrations. b) Schematic model illustrating the effect of increasing K_D_ on the acceptor FRET response predicted by a quadratic binding model. c) Donor fluorescence quenching of mScarlet3–SpyTag003 upon titration with AF647–SpyCatcher003 at varying donor concentrations. d) Schematic model showing the impact of increasing K_D_ on donor quenching behavior in a quadratic model. e) Comparison of K_D_ determined for mScarlet3–SpyTag003 binding to AF647–SpyCatcher003 under different assay conditions. Bars represent mean values from three independently prepared replicates; error bars indicate the standard error.

However, this approach is highly sensitive to accurate correction of spectral bleed-through, including donor emission leaking into the acceptor channel and direct excitation of the acceptor at the donor excitation wavelength, both of which generate non-FRET signals.^44^

In the donor-quenching approach, the donor fluorescence is monitored during the same titration series. Different concentrations of donor protein are mixed with varying amounts of acceptor protein, and a quadratic binding model is fit to the resulting quenching curves in the same manner as for the acceptor-emission method (Figure 3c). As with acceptor-emission analysis, tighter binders with lower K_D_ values produce steeper, more cooperative-looking quenching curves (Figure 3d). In this approach, the only bleed-through signal is the fluorescence of the acceptors in the donor emission, which can be calculated and subtracted which makes the calculations simpler, especially when reactions are performed in more complex environments such as crude lysates.

To evaluate the accuracy of HTFAST for measuring K_D_, we quantified the affinity between SpyTag003 and SpyCatcher003 using both crude lysate and purified protein samples. The purified mScarlet3–SpyTag003 fusion yielded a K_D_ of 3.2 ± 0.3 nM by acceptor-emission analysis, closely matching the previously reported value of ∼3 nM obtained using a stopped-flow FRET assay.^45^In crude lysates, both donor-quenching,

3.5 ± 1.4 nM and acceptor-emission 4.5 ± 1.4 nM, methods produced higher sample-to-sample variability; however, the resulting K_D_ values were not statistically different from those obtained with purified protein.

### Nanobody and sdAbs screening using HTFAST

To evaluate the capability of HTFAST for nanobody screening, we tested a previously characterized nanobody targeting the human CD4 surface protein, Nb457. Nb457 binds CD4 with high affinity, exhibiting a reported K_D_ of 4.2 nM for the Nb457–NbHSA–Nb457 complex (PDB: 8W90).^46^The mScarlet3–Nb457 fusion was expressed in the CFE system and used as the donor molecule, while commercial human CD4 protein was conjugated to AF647 to generate the acceptor (Figure 4a). Crude, unpurified donor protein was titrated with the labeled CD4 acceptor, and K_D_ values were determined using both donor-quenching and acceptor-emission analyses. The donor-quenching method yielded more consistent measurements and superior model fit (Supporting information Figure S3 and S4). For the acceptor emission method K_D_ of 27.4 ± 16.1 nM was observed (**Figure 4b**). The K_D_ of 3.9 ± 0.6 nM for Nb457 binding to human CD4 measured with Donor-quenching is compatible with K_D_ of 4.2 nM for the Nb457– NbHSA–Nb457 complex previously reported.^46^It should be noted that when working with crude samples, the donor-quenching method is superior due to its simplicity in both measurement and analysis, as it does not require bleed-through correction factors such as α and β. This approach is also more robust in turbid solutions, such as crude CFE lysates.

**Figure 4.**
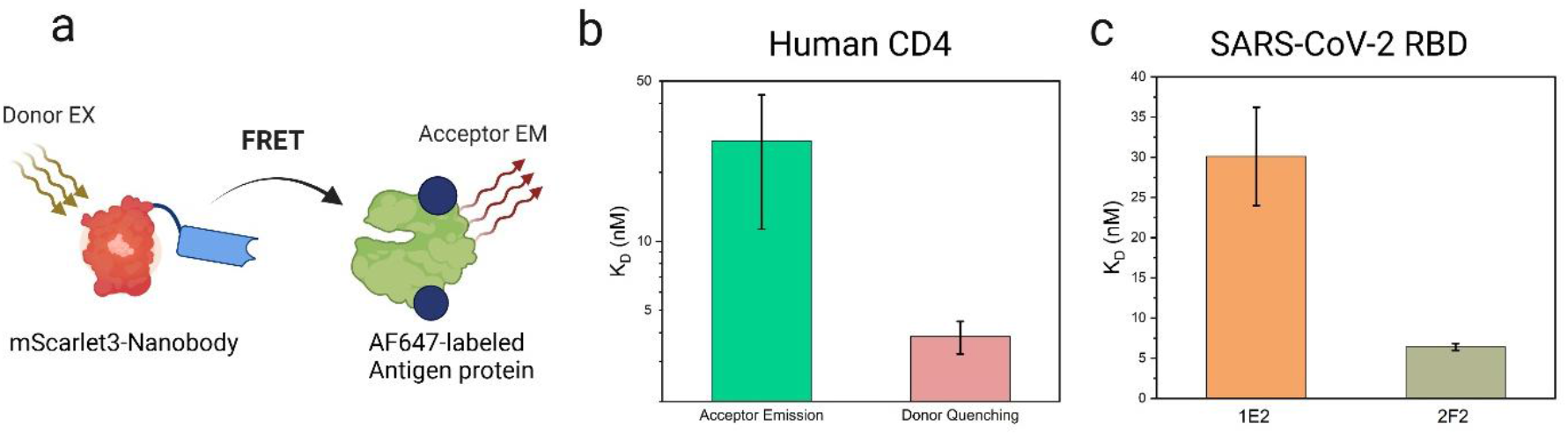
a) Schematic of the FRET interaction between the mScarlet3–nanobody fusion (donor) and its AF647-labeled antigen (acceptor). b) Equilibrium dissociation constant K_D_ of Nb457 binding to AF647-labeled human CD4. The green bar represents the K_D_ determined from acceptor-emission analysis, and the red bar represents the K_D_ obtained from donor-quenching analysis. Bars show the mean of three independent replicates; error bars represent the standard error. c) Equilibrium dissociation constants of different sdAbs binding to AF647-labeled SARS-CoV-2 RBD. Bars show the mean of three independent replicates; error bars represent the standard error.

We further evaluated the capability of HTFAST to rank protein binders targeting the same antigen. The SARS-CoV-2 receptor-binding domain (RBD) has been extensively characterized, and a diverse panel of single-domain antibodies (sdAbs) with a wide range of reported K_D_ values has been published.^47^ From this panel, we selected three sdAbs with reported K_D_s, spanning low-to mid-nanomolar affinities, 1E2 (K_D_ = 35.5 nM), 2F2 (K_D_ = 5.18 nM), and 5F8 (K_D_ = 0.996 nM) to benchmark HTFAST’s ability to resolve affinity differences that could be observed in a DBTL cycle study.

mScarlet3-fused sdAbs were expressed in the CFE system and titrated against AF647-labeled SARS-CoV-2 RBD. Using the donor-quenching analysis, 1E2 yielded a K_D_ of 30.1 ± 6.1 nM while 2F2 exhibited a tighter binder at 6.4 ± 0.8 nM consistent with their reported relative strengths (**Figure 4c**). In contrast, 5F8 produced no measurable FRET signal despite exhibiting robust donor fluorescence, indicating successful expression of the fusion protein but suggesting that it was nonfunctional in our tests. This could be due to the differences in the protein expression systems used, which can lead to different properties of the final folded protein.

### Expression of both binder and ligand in CFE systems

To further increase the throughput of binder discovery, we evaluated whether both the binder and ligand could be co-expressed and quantified directly within the cell-free expression system. In this configuration, fluorescent proteins were fused to SpyTag003 and SpyCatcher003 to generate donor–acceptor FRET pairs. We retained the previously optimized mScarlet3–SpyTag003 construct and selected mCitrine (Ex 516 nm, Em 529 nm) to label SpyCatcher003. In earlier experiments using Alexa Fluor 647, mScarlet3 functioned as the donor; however, in this new fluorescent-protein pair, mCitrine serves as the donor and mScarlet3 as the acceptor. Using this configuration, we observed that donor-quenching analysis was not feasible, whereas acceptor-emission measurements produced robust FRET signals. From these measurements, we determined an apparent K_D_ of 4.9 ± 1.3 nM (mean ± SE, n = 3 independent replicates).

It should be noted that the same mCitrine–mScarlet3 fluorescent-protein pair was used to detect FRET between the nanobody Nb6101 and the BCL11A ZF456 protein, which were previously reported to form a high-affinity complex.^48^However, in our CFE-expressed system, a poor FRET signal was observed between these two binders (Supplement S5). Also, we tried expressing and purifying ZF456 to further test the Nb6101 which we couldn’t get a proper ZF456 protein (See supplement section 4 for detailed method). We additionally replaced mCitrine with another fluorescent protein, mTurquoise2 (Ex 434 nm, Em 474 nm), to modify the FRET pair geometry; however, this substitution did not improve the FRET signal. Notably, blue and cyan fluorescent proteins were evaluated twice in this study, and both attempts were unsuccessful. This outcome may be attributed to interference from *E. coli* cell-free system autofluorescence within the blue–cyan spectral region.

## Conclusion

In summary, we have developed and optimized the High-Throughput FRET Affinity Screening Technology (HTFAST) for rapid screening and characterization of kinetic binding parameters of nanobodies, single-domain antibodies (sdAbs), and purification tags in cell-free expression systems, without requiring purification of the expressed proteins. This approach can be readily integrated into design–build–test–learn (DBTL) cycles in cell-free platforms to accelerate the engineering of improved binding proteins. However, more detailed kinetic analyses, providing additional parameters such as association (k_on_) and dissociation (k_off_) rates, should be performed after the top binders are selected. It should also be noted that, due to the distance dependency of FRET, larger proteins such as full-length antibodies may require further optimization or alternative assay designs to observe a robust FRET response.

## Funding

Research reported in this publication was supported by NIGMS of the National Institutes of Health under award number R35GM138265. The content is solely the responsibility of the authors and does not necessarily represent the official views of the National Institutes of Health. This work was also supported in part by NSF Award #: 2242763.

## Acknowledgments

We thank Dr. Scott Nelson, Assistant Professor at Iowa State University, for his invaluable guidance and support with the FRET measurements and modeling analyses. We thank Ryan Godin for useful discussions on cell-free expression techniques.

## Notes

### Competing Interest Statement

The authors have declared no competing interest.

## References

1. Lu, R.-M. et al. Development of therapeutic antibodies for the treatment of diseases. J Biomed Sci 27, 1 (2020).

2. Liu, J. et al. Unveiling the new chapter in nanobody engineering: advances in traditional construction and AI-driven optimization. J Nanobiotechnol 23, 87 (2025).

3. Harmsen, M. M. & De Haard, H. J. Properties, production, and applications of camelid single-domain antibody fragments. Appl Microbiol Biotechnol 77, 13–22 (2007).

4. Jovčevska, I. & Muyldermans, S. The Therapeutic Potential of Nanobodies. BioDrugs 34, 11–26 (2020).

5. Zhu, L. et al. Highly potent and broadly neutralizing anti-CD4 trimeric nanobodies inhibit HIV-1 infection by inducing CD4 conformational alteration. Nat Commun 15, 6961 (2024).

6. Van Der Linden, R. H. J. et al. Comparison of physical chemical properties of llama VHH antibody fragments and mouse monoclonal antibodies. Biochimica et Biophysica Acta (BBA) - Protein Structure and Molecular Enzymology 1431, 37–46 (1999).

7. Yin, M. et al. Evolution of nanobodies specific for BCL11A. Proc. Natl. Acad. Sci. U.S.A. 120, e2218959120 (2023).

8. Zimmermann, I. et al. Synthetic single domain antibodies for the conformational trapping of membrane proteins. eLife 7, e34317 (2018).

9. Miton, C. M. & Tokuriki, N. How mutational epistasis impairs predictability in protein evolution and design. Protein Science 25, 1260–1272 (2016).

10. Meger, A. T. et al. Rugged fitness landscapes minimize promiscuity in the evolution of transcriptional repressors. Cell Systems 15, 374-387.e6 (2024).

11. Landwehr, G. M. et al. Accelerated enzyme engineering by machine-learning guided cell-free expression. Nat Commun 16, 865 (2025).

12. Thornton, E. L., Boyle, J. T., Laohakunakorn, N. & Regan, L. Cell-Free Protein Synthesis as a Method to Rapidly Screen Machine Learning-Generated Protease Variants. ACS Synth. Biol. 14, 1710–1718 (2025).

13. Thornton, E. L. et al. Applications of cell free protein synthesis in protein design. Protein Science 33, e5148 (2024).

14. Ojima-Kato, T., Nagai, S. & Nakano, H. Ecobody technology: rapid monoclonal antibody screening method from single B cells using cell-free protein synthesis for antigen-binding fragment formation. Sci Rep 7, 13979 (2017).

15. Ojima-Kato, T. et al. ‘Zipbody’ leucine zipper-fused Fab in E. coli in vitro and in vivo expression systems. Protein Engineering, Design and Selection 29, 149–157 (2016).

16. Chen, X., Gentili, M., Hacohen, N. & Regev, A. A cell-free nanobody engineering platform rapidly generates SARS-CoV-2 neutralizing nanobodies. Nat Commun 12, 5506 (2021).

17. Hunt, A. C. et al. A rapid cell-free expression and screening platform for antibody discovery. Nat Commun 14, 3897 (2023).

18. Beaudet, L. et al. AlphaLISA immunoassays: the no-wash alternative to ELISAs for research and drug discovery. Nat Methods 5, an8–an9 (2008).

19. Matsunaga, R. et al. High-throughput analysis system of interaction kinetics for data-driven antibody design. Sci Rep 13, 19417 (2023).

20. Dzimianski, J. V. et al. Rapid and sensitive detection of SARS-CoV-2 antibodies by biolayer interferometry. Sci Rep 10, 21738 (2020).

21. Yasgar, A., Jadhav, A., Simeonov, A. & Coussens, N. P. AlphaScreen-Based Assays: Ultra-High-Throughput Screening for Small-Molecule Inhibitors of Challenging Enzymes and Protein-Protein Interactions. in High Throughput Screening (ed. Janzen, W. P.) vol. 1439 77–98 (Springer New York, New York, NY, 2016).

22. Hunt, A. C. et al. A rapid cell-free expression and screening platform for antibody discovery. Nat Commun 14, 3897 (2023).

23. Eglen, R. M. et al. The Use of AlphaScreen Technology in HTS: Current Status. Curr Chem Genomics 1, 2–10 (2008).

24. Margineanu, A. et al. Screening for protein-protein interactions using Förster resonance energy transfer (FRET) and fluorescence lifetime imaging microscopy (FLIM). Sci Rep 6, 28186 (2016).

25. Basu, S. et al. FRET-enhanced photostability allows improved single-molecule tracking of proteins and protein complexes in live mammalian cells. Nat Commun 9, 2520 (2018).

26. Dopp, J. L., Jo, Y. R. & Reuel, N. F. Methods to reduce variability in E. Coli-based cell-free protein expression experiments. Synthetic and Systems Biotechnology 4, 204–211 (2019).

27. Hejazi, S., Godin, R., Jurasic, V. & Reuel, N. F. Single-Walled Carbon Nanotube Probes for Protease Characterization Directly in Cell-Free Expression Reactions. Anal. Chem. 97, 10745–10754 (2025).

28. Hejazi, S., Ahsan, A., Kashani, S., Tameiv, D. & Reuel, N. F. Amplified DNA heterogeneity assessment with Oxford Nanopore sequencing applied to cell free expression templates. PLoS ONE 19, e0305457 (2024).

29. Dopp, J. L. & Reuel, N. F. Simple, functional, inexpensive cell extract for in vitro prototyping of proteins with disulfide bonds. Biochemical Engineering Journal 164, 107790 (2020).

30. Godin, R., Hejazi, S., Lange, B., Aldamak, B. & Reuel, N. F. Rapid Cell-Free Combinatorial Mutagenesis Workflow Using Small Oligos Suitable for High-Iteration, Active Learning-Guided Protein Engineering. Preprint at 10.1101/2025.06.18.660386 (2025).

31. Weitner, T., Friganović, T. & Šakić, D. Inner Filter Effect Correction for Fluorescence Measurements in Microplates Using Variable Vertical Axis Focus. Anal. Chem. 94, 7107–7114 (2022).

32. Liao, J., Madahar, V., Dang, R. & Jiang, L. Quantitative FRET (qFRET) Technology for the Determination of Protein–Protein Interaction Affinity in Solution. Molecules 26, 6339 (2021).

33. Chen, H., Puhl, H. L., Koushik, S. V., Vogel, S. S. & Ikeda, S. R. Measurement of FRET Efficiency and Ratio of Donor to Acceptor Concentration in Living Cells. Biophysical Journal 91, L39–L41 (2006).

34. Willi, J. A., Karim, A. S. & Jewett, M. C. Cell-Free Translation Quantification via a Fluorescent Minihelix. ACS Synth. Biol. 13, 2253–2259 (2024).

35. Keeble, A. H. et al. Approaching infinite affinity through engineering of peptide– protein interaction. Proc. Natl. Acad. Sci. U.S.A. 116, 26523–26533 (2019).

36. Surre, J. et al. Strong increase in the autofluorescence of cells signals struggle for survival. Sci Rep 8, 12088 (2018).

37. Mihalcescu, I., Van-Melle Gateau, M., Chelli, B., Pinel, C. & Ravanat, J.-L. Green autofluorescence, a double edged monitoring tool for bacterial growth and activity in micro-plates. Phys. Biol. 12, 066016 (2015).

38. Gadella, T. W. J. et al. mScarlet3: a brilliant and fast-maturing red fluorescent protein. Nat Methods 20, 541–545 (2023).

39. Legault, S. et al. Generation of bright monomeric red fluorescent proteins via computational design of enhanced chromophore packing. Chem. Sci. 13, 1408–1418 (2022).

40. Stoddard, A. & Rolland, V. I see the light! Fluorescent proteins suitable for cell wall/apoplast targeting in Nicotiana benthamiana leaves. Plant Direct 3, e00112 (2019).

41. Mo, G. C. H., Posner, C., Rodriguez, E. A., Sun, T. & Zhang, J. A rationally enhanced red fluorescent protein expands the utility of FRET biosensors. Nat Commun 11, 1848 (2020).

42. Chu, J. et al. A bright cyan-excitable orange fluorescent protein facilitates dual-emission microscopy and enhances bioluminescence imaging in vivo. Nat Biotechnol 34, 760–767 (2016).

43. Lambert, T. J. FPbase: a community-editable fluorescent protein database. Nat Methods 16, 277–278 (2019).

44. Jiang, L. et al. Protein–Protein Affinity Determination by Quantitative FRET Quenching. Sci Rep 9, 2050 (2019).

45. Keeble, A. H. et al. Approaching infinite affinity through engineering of peptide– protein interaction. Proc. Natl. Acad. Sci. U.S.A. 116, 26523–26533 (2019).

46. Zhu, L. et al. Highly potent and broadly neutralizing anti-CD4 trimeric nanobodies inhibit HIV-1 infection by inducing CD4 conformational alteration. Nat Commun 15, 6961 (2024).

47. Chi, X. et al. Humanized single domain antibodies neutralize SARS-CoV-2 by targeting the spike receptor binding domain. Nat Commun 11, 4528 (2020).

48. Yin, M. et al. Evolution of nanobodies specific for BCL11A. Proc. Natl. Acad. Sci. U.S.A. 120, e2218959120 (2023).

