## Supporting information for "High-Throughput FRET Affinity Screening Technique (HTFAST) For Cell-Free Expressed Binding Protein Characterization"

### Section 1: Calibration curves for FL protein quantification

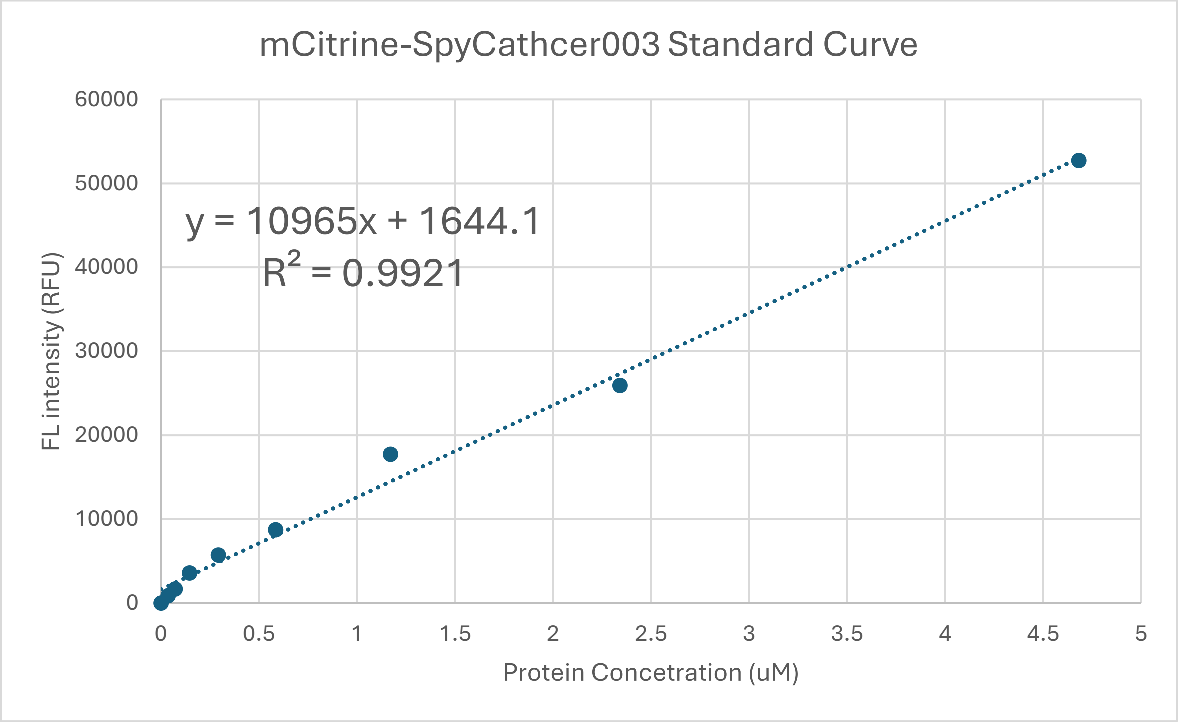

*Figure S1 mCitrine-Spycatcher003 concentration calibration curve. Known amounts purified fluorescent protein were added to cell-free reaction mixtures at defined dilution ratios, and the fluorescent signal of each was measured. The points present the average of three measurements, and the error bars are the standard error.*

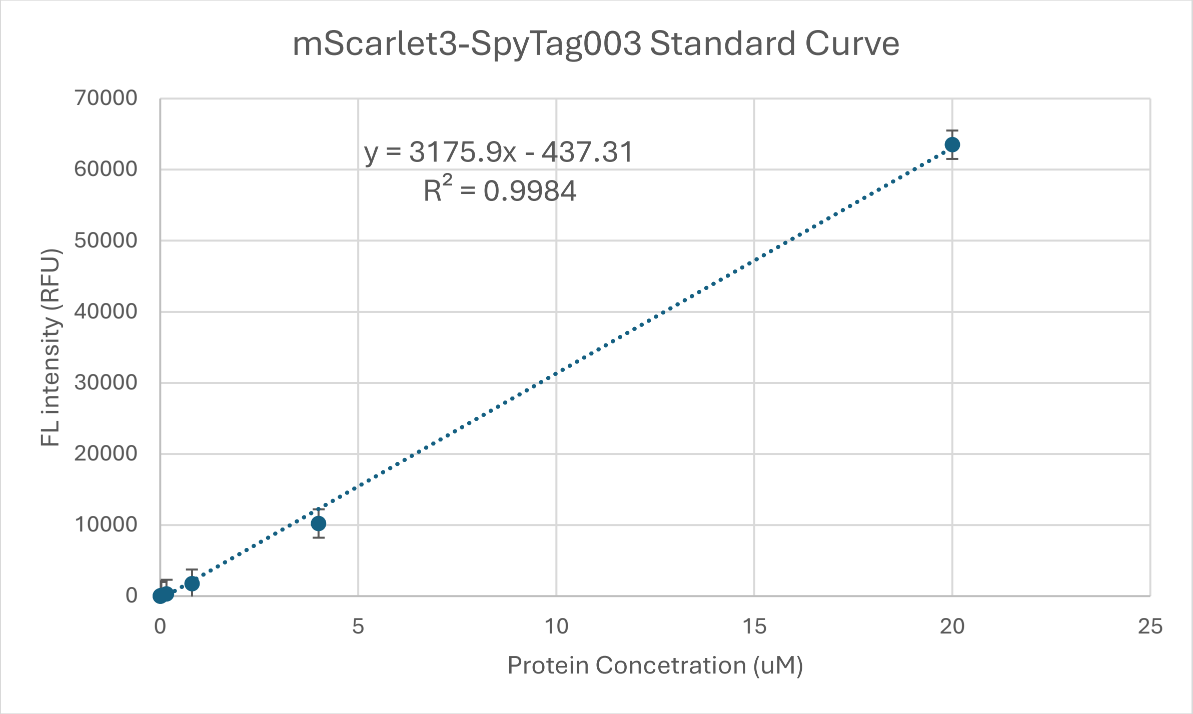

*Figure S2 mScarlet-SpyTag003 concentration calibration curve. Known amounts purified fluorescent protein were added to cell-free reaction mixtures at defined dilution ratios, and the fluorescent signal of each was measured. The points present the average of three measurements, and the error bars are the standard error.*

### Section 2: Derivation of Acceptor Emission Equation

We assume a simple 1:1 binding equilibrium between a donor-labeled species $D$and an acceptor-labeled species $A$:

$$D + A \rightleftharpoons DA$$

with dissociation constant

$$K_d = \frac{[D][A]}{[DA]}.$$

Let the *total* donor and acceptor concentrations be

$$D_T = [D] + [DA], A_T = [A] + [DA].$$

Defining $x = [DA]$, we have

$$[D] = D_T - x, [A] = A_T - x.$$

Substituting into the definition of $K_d$:

$$K_d = \frac{(D_T - x)(A_T - x)}{x}.$$

Rearranging gives a quadratic equation in $x$:

$$xK_d = (D_T - x)(A_T - x) = D_T A_T - x(D_T + A_T) + x^2,$$
$$x^2 - x(D_T + A_T + K_d) + D_T A_T = 0.$$

Solving this quadratic for $x = [DA]$and choosing the physically meaningful (smaller) root:

$$[DA] = \frac{D_T + A_T + K_d - \sqrt{(D_T + A_T + K_d)^2 - 4D_T A_T}}{2}.$$

The FRET signal is assumed to be proportional to the fraction of donor molecules that are bound to acceptor. Thus, if $E_{\text{FRET,max}}$is the FRET emission when all donor is bound, the observed FRET emission is

$$E_{\text{FRET}} = E_{\text{FRET,max}} f_{\text{bound}},$$

where

$$f_{\text{bound}} = \frac{[DA]}{D_T}.$$

Substituting the expression for $[DA]$and simplifying yields

$$f_{\text{bound}} = 1 - \frac{2K_d}{A_T - D_T + K_d + \sqrt{(A_T - D_T - K_d)^2 + 4K_dA_T}}.$$

Therefore,

$$E_{\text{FRET}} = E_{\text{FRET,max}}\left(1 - \frac{2K_d}{A_T - D_T + K_d + \sqrt{(A_T - D_T - K_d)^2 + 4K_dA_T}}\right),$$

which is the form used in this work to fit the FRET titration curves.

**Derivation of Donor Quenching Equation:**

The fraction of donor molecules that are bound to acceptor is

$$f_{\text{bound}} = \frac{[DA]}{D_T} = \frac{D_T + A_T + K_d - \sqrt{(D_T + A_T + K_d)^2 - 4D_TA_T}}{2D_T}.$$

In the donor channel (DD), the fluorescence intensity decreases as more donors are bound and quenched by FRET. We assume a linear relationship between donor quenching and the fraction of donors in the bound state:

- $FL_{DD}^{0}$: donor-channel fluorescence when no acceptor is bound (all donors unquenched, $f_{\text{bound}} = 0$).
- $FL_{DD}^{\min}$: donor-channel fluorescence when all donors are bound and maximally quenched ($f_{\text{bound}} = 1$).

Thus, the observed donor-channel fluorescence is

$$FL_{DD} = FL_{DD}^{0} - (FL_{DD}^{0} - FL_{DD}^{\min})f_{\text{bound}}.$$

Substituting the expression for $f_{\text{bound}}$gives

$$FL_{DD} = FL_{DD}^{0} - (FL_{DD}^{0} - FL_{DD}^{\min})\left(\frac{D_T + A_T + K_d - \sqrt{(D_T + A_T + K_d)^2 - 4D_TA_T}}{2D_T}\right),$$

which can be written as

$$FL_{DD} = FL_{DD}^{0} - (FL_{DD}^{0} - FL_{DD}^{\min})\frac{D_T + A_T + K_d - \sqrt{(D_T + A_T + K_d)^2 - 4D_TA_T}}{2D_T}$$

the form used in this work to fit donor-quenching binding curves.

### Section 3: Fitting of FRET data to the quadratic models using Acceptor emission and Donor quenching.

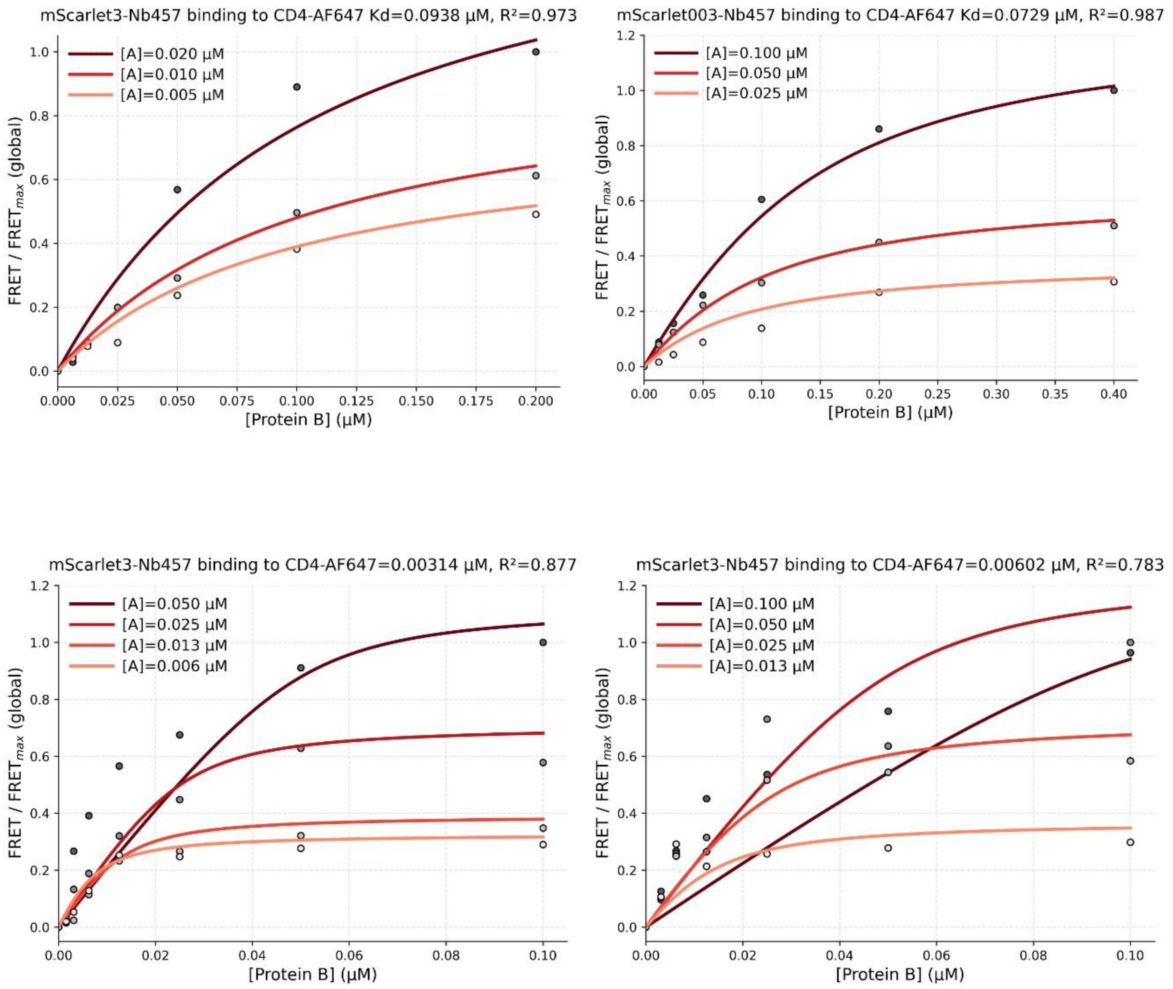

***Figure S3.*** *Acceptor emission (FRET) response of mScarlet3–Nb457 (donor, Protein A) upon titration with CD4–AF647 (acceptor, Protein B) at varying donor concentrations. Each panel represents an independent experimental replicate generated from separate cell-free expression (CFE) reactions performed on different days.*

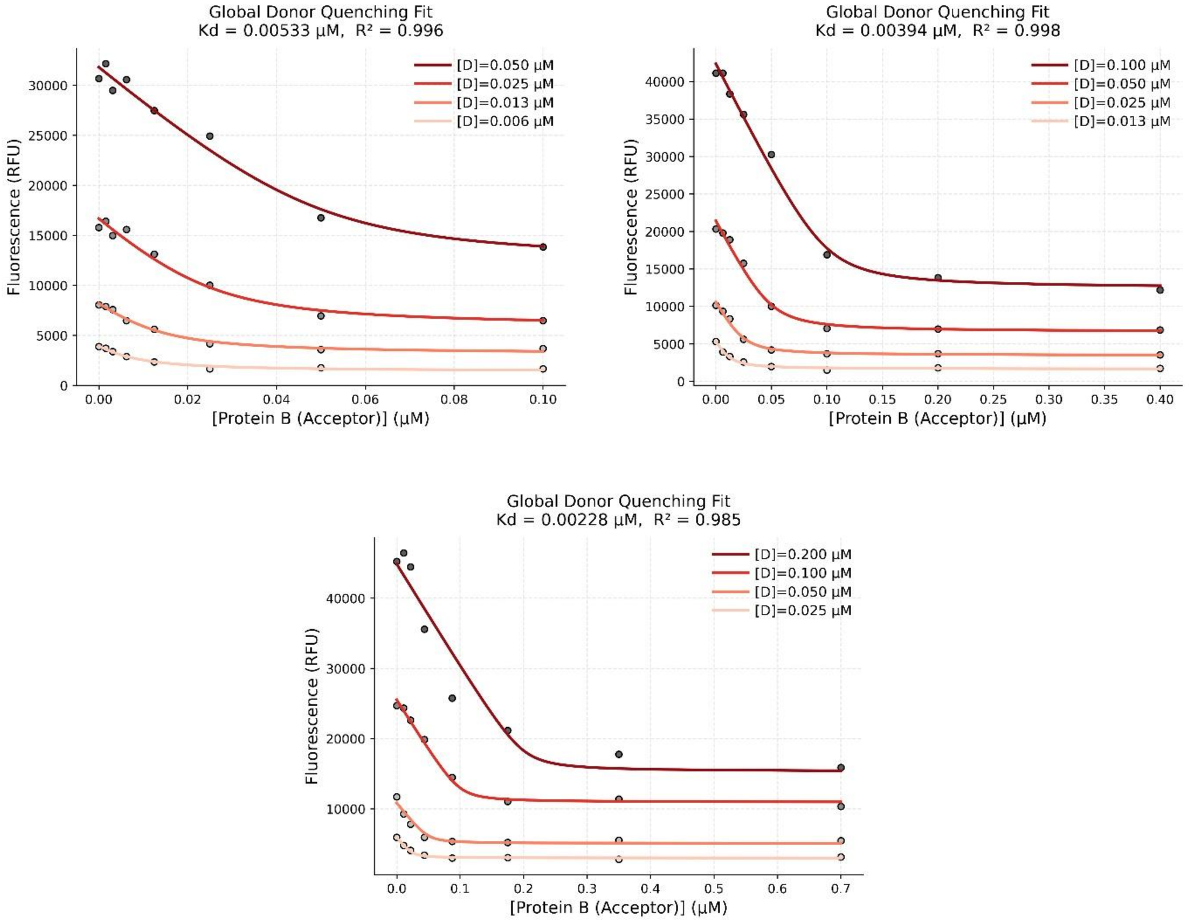

***Figure S4 .*** *Donor quenching response of mScarlet3–Nb457 (donor, Protein A) upon titration with CD4–AF647 (acceptor, Protein B) at varying donor concentrations. Donor fluorescence decreases with increasing acceptor concentration due to FRET. Each panel represents an independent experimental replicate generated from separate cell-free expression (CFE) reactions performed on different days.*

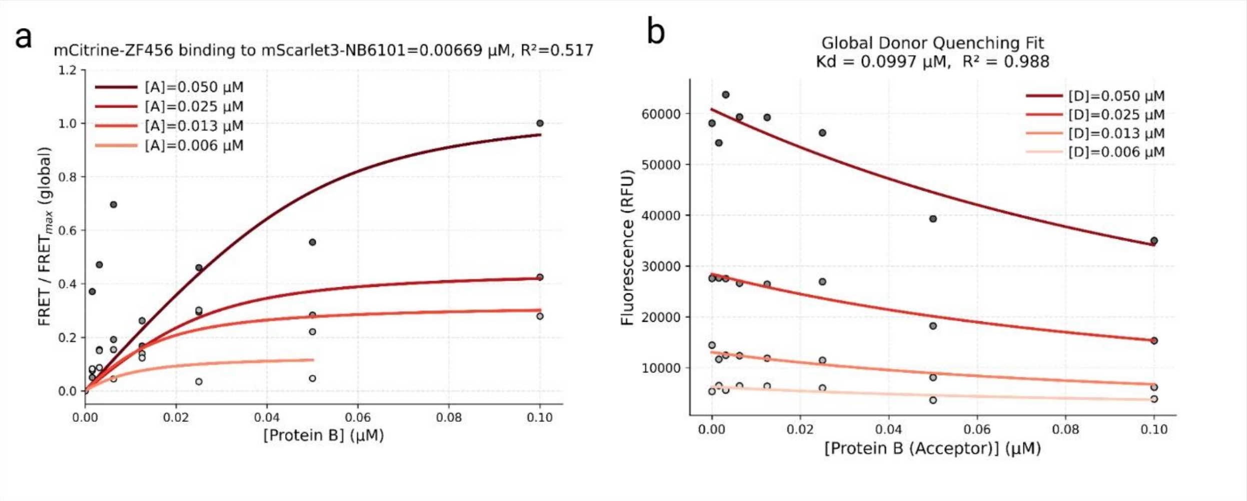

*Figure S5 a) acceptor emission response of mCitrine-ZF456 (donor, Protein A) upon titration with mScarlet-Nb6101 (acceptor, Protein B) at varying donor concentrations. Donor fluorescence decreases with increasing acceptor concentration due to FRET. b) Donor quenching response of mCitrine-ZF456 (donor, Protein A) upon titration with mScarlet-Nb6101 (acceptor, Protein B) at varying donor concentrations. Donor fluorescence decreases with increasing acceptor concentration due to FRET.*

### Section 4: Cloning and Protein Purification.

Method:

DNA sequence encoding ZF456 part of BCL11A (residues 737 to 835) was optimized and synthesized through IDT. This insert was amplified using primers with 30bp overlapping sequences on either side of the insertion site. The final insert was cloned into a pET26b vector using Gibson assembly at the BamHI site, following the supplier's protocol (New England Biolabs, cat # E2611). The assembled product was transformed into freshly prepared electrocompetent E. coli BL21 (Tu et al. 2016). The cells were then recovered in LB medium for one hour and plated on LB with Kanamycin (50 μg/mL) overnight. Positive colonies were confirmed through colony PCR and sequencing at Iowa State University's DNA Facility. Escherichia coli BL21 cells harboring the plasmid were cultured in LB medium supplemented with kanamycin (50 μg/mL) and glucose (0.1 M) at 37 °C with shaking at 270 rpm. When the culture reached an $OD_{600}$ of 0.6, protein expression was induced by the addition of IPTG (1 mM). Following induction, the culture was cooled to 18 °C and incubated overnight with shaking at 270 rpm.

Cells harvested following our previous protocol.[1] The cells were lysed, and the recombinant proteins were purified using a Strep-Tactin® XT 4Flow® high-capacity column (IBA Lifesciences). Mass spectrometry analysis of the purified proteins was performed at the Iowa State University Protein Facility. Based on the protein sequence, a molecular weight of 12.63 kDa was expected; however, the mass spectrometry data revealed peaks at 12.22 kDa and 16.91 kDa.

### Section 5: DNA templates used in this study.

Spytag3-super-TagRFP:

GTAAAACGACGGCCAGTAGCGCTATTAAAGCTTCGAAATTAATACGACTCACTATA
GGGAGACCACAACGGTTTCCCTCTAGAAATAATTTTGTTTAACTTTAAGAAGGAGAT
ATACATATGGGACGTGGCGTTCCTCATATTGTTATGGTGGACGCCTACAAACGCTA
TAAATCGGGAGGTGGTTCAGGCATGGTATCTAAAGGGGAGGAACTTATTAAAGAAA
ACATGCACATGAAATTGTACATGGAGGGCACTGTCAATAACCACCACTTCAAGTGT
ACTTCTGAGGGGGAAGGTAAGCCGTATGAGGGTACGCAAACCATGCGCATCAAGG
TGGTTGAGGGGGGTCCACTCCCATTTGCGTTTGACATTCTGGCGACGAGTTTTATG
TATGGGTCGCGTACCTTCATTAATCACACCCAGGGTATCCCGGACTTCTTTAAACA
ATCGTTTCCGGAAGGCTTCACATGGGAGCGTGTTACCACTTACGAGGACGGGGGG
GTATTAACCGCGACCCAAGACACATCCTTACAAGATGGCTGTCTGATTTATAATGT
GAAAATTCGCGGTGTTAATTTCCCATCCAACGGTCCTGTTATGCAGAAAAAGACAC
TGGGGTGGGAGGCCAACACCGAAATGTTGTATCCGGCCGACGGGGGTTTAGAGG
GGCGCACAGTAATGGCCCTTAAACTGGTCGGTGGCGGCCACTTAATCTGCAATTT
CAAGACAACGTATCGGTCTAAGAAACCTGCCAAAAACCTTAAGATGCCAGGGGTAT
ATTACGTGGACCATCGGTTAGAACGGATTAAGGAGGCTGATAAGGAGACTTACGTT
GAGCAGCATGAAGTGGCGGTAGCGCGGTATTGCGACCTGCCTTCCAAGTTGGGG
CACAAGCTCAACGGGATGGACGAGTTGTACAAGTGGAGCCATCCGCAGTTCGAAA
AATAATAAGTCGACCGGCTGCTAACAAAGCCCGAAAGGAAGCTGAGTTGGCTGCT
GCCACCGCTGAGCAATAACTAGCATAACCCCTTGGGGCCTCTAAACGGGTCTTGA
GGGGTTTTTTGCTGAAAGCGAGACTAAGCTTTAAACTTCGGGTCATAGCTGTTTCC
TG

Spytag3-mKate2:

GTAAAACGACGGCCAGTAGCGCTATTAAAGCTTCGAAATTAATACGACTCACTATA
GGGAGACCACAACGGTTTCCCTCTAGAAATAATTTTGTTTAACTTTAAGAAGGAGAT
ATACATATGGGACGTGGCGTTCCTCATATTGTTATGGTGGACGCCTACAAACGCTA
TAAATCGGGAGGTGGTTCAGGCATGGTGAGTGAACTCATCAAGGAGAACATGCAT
ATGAAATTATACATGGAGGGGACGGTCAACAATCATCACTTCAAGTGTACCTCGGA
AGGTGAAGGTAAACCATATGAGGGGACGCAGACTATGCGTATCAAGGCGGTTGAG
GGCGGCCCACTTCCATTCGCATTTGACATCTTAGCTACATCCTTTATGTATGGCTC
CAAGACATTCATTAATCATACACAGGGCATCCCAGACTTCTTTAAGCAGTCGTTCC
CTGAAGGGTTCACCTGGGAACGTGTCACGACATATGAAGACGGGGGCGTCCTTAC
GGCAACTCAGGACACGTCGTTGCAAGATGGTTGCCTGATCTATAACGTTAAGATCC
GCGGGGTGAATTTCCCTTCTAATGGGCCTGTAATGCAAAAGAAGACATTAGGCTG
GGAGGCCAGTACAGAGACCCTTTATCCGGCCGATGGTGGCTTGGAAGGTCGCGC
GGACATGGCCCTGAAATTGGTCGGGGGCGGCCACTTGATCTGCAATCTGAAGACG
ACTTACCGTAGCAAGAAGCCAGCTAAGAATCTCAAAATGCCGGGTGTATATTATGT
AGACCGCCGCTTGGAGCGGATCAAGGAGGCAGACAAAGAAACGTATGTAGAACAA
CATGAAGTCGCAGTCGCTCGCTATTGTGACTTGCCATCAAAATTGGGCCATCGCTG
GAGCCATCCGCAGTTCGAAAAATAATAAGTCGACCGGCTGCTAACAAAGCCCGAA

AGGAAGCTGAGTTGGCTGCTGCCACCGCTGAGCAATAACTAGCATAACCCCTTGG
GGCCTCTAAACGGGTCTTGAGGGGTTTTTTGCTGAAAGCGAGACTAAGCTTTAAAC
TTCGGGTCATAGCTGTTTCCTG

Spytag3-TagRFP:

GTAAAACGACGGCCAGTAGCGCTATTAAAGCTTCGAAATTAATACGACTCACTATA
GGGAGACCACAACGGTTTCCCTCTAGAAATAATTTTGTTTAACTTTAAGAAGGAGAT
ATACATATGGGACGTGGCGTTCCTCATATTGTTATGGTGGACGCCTACAAACGCTA
TAAATCGGGAGGTGGTTCAGGCATGTCGGAGCTGATTAAAGAAAACATGCATATGA
AGTTGTACATGGAAGGCACGGTAAACAACCATCATTTCAAGTGTACATCGGAGGGT
GAGGGTAAACCATACGAGGGTACGCAAACCATGCGTATCAAGGTTGTCGAGGGGG
GCCCGCTCCCGTTCGCGTTTGATATTCTGGCAACTTCTTTTATGTACGGTTCGCGC
ACTTTCATTAACCACACTCAAGGCATTCCAGATTTTTTCAAGCAATCATTCCCTGAG
GGGTTTACGTGGGAACGTGTCACCACTTATGAGGACGGTGGCGTCCTGACAGCTA
CTCAAGACACATCACTCCAGGATGGCTGTCTTATCTATAATGTGAAGATTCGGGGC
GTTAACTTTCCGTCGAACGGGCCTGTAATGCAAAAAAAGACCCTTGGGTGGGAAG
CTAATACTGAAATGTTGTACCCAGCGGACGGCGGGCTCGAAGGGCGGAGTGACAT
GGCATTGAAATTGGTAGGCGGTGGCCACCTTATTTGCAATTTCAAGACGACGTACC
GCTCCAAGAAACCTGCTAAGAACCTGAAGATGCCAGGCGTCTATTACGTTGACCAC
CGCCTTGAACGGATTAAAGAAGCAGACAAAGAGACCTACGTTGAACAGCATGAGG
TCGCAGTTGCCCGCTATTGTGATCTGCCTTCAAAACTCGGTCACAAATGGAGCCAT
CCGCAGTTCGAAAAATAATAAGTCGACCGGCTGCTAACAAAGCCCGAAAGGAAGC
TGAGTTGGCTGCTGCCACCGCTGAGCAATAACTAGCATAACCCCTTGGGGCCTCT
AAACGGGTCTTGAGGGGTTTTTTGCTGAAAGCGAGACTAAGCTTTAAACTTCGGGT
CATAGCTGTTTCCTG

Spytag3-CyOFP1:

GTAAAACGACGGCCAGTAGCGCTATTAAAGCTTCGAAATTAATACGACTCACTATA
GGGAGACCACAACGGTTTCCCTCTAGAAATAATTTTGTTTAACTTTAAGAAGGAGAT
ATACATATGGGACGTGGCGTTCCTCATATTGTTATGGTGGACGCCTACAAACGCTA
TAAATCGGGAGGTGGTTCAGGCATGGTAAGTAAGGGTGAGGAGCTGATTAAAGAG
AATATGCGCTCCAAACTCTACTTGGAGGGCAGTGTGAACGGCCACCAGTTTAAGT
GCACCCATGAGGGGGAAGGCAAACCTTACGAAGGTAAACAGACGAACCGTATCAA
GGTGGTCGAAGGCGGGCCTTTGCCATTCGCATTCGACATCTTAGCCACCCATTTCA
TGTATGGCAGTAAAGTGTTCATTAAATACCCGGCTGATCTCCCGGATTATTTTAAAC
AATCGTTCCCAGAGGGGTTTACCTGGGAACGGGTGATGGTTTTCGAGGACGGTGG
TGTTTTGACGGCTACTCAAGATACATCTCTGCAAGATGGTGAACTTATTTACAACGT
CAAAGTACGGGGGGTAAACTTTCCAGCCAACGGTCCTGTTATGCAGAAGAAGACT
TTGGGTTGGGAACCTTCCACAGAAACAATGTATCCGGCGGATGGCGGGCTGGAG
GGGCGCTGCGATAAAGCTCTTAAATTGGTTGGCGGTGGGCATCTTCATGTTAATTT

CAAAACAACATACAAATCTAAGAAGCCGGTTAAAATGCCAGGGGTCCACTACGTTG
ACCGCCGCTTAGAGCGTATCAAAGAAGCAGACAATGAGACTTATGTGGAACAGTA
CGAACATGCAGTCGCCCGGTACTCAAACCTCGGTGGCGGCATGGATGAACTGTAC
AAGTGGAGCCATCCGCAGTTCGAAAAATAATAAGTCGACCGGCTGCTAACAAAGC
CCGAAAGGAAGCTGAGTTGGCTGCTGCCACCGCTGAGCAATAACTAGCATAACCC
CTTGGGGCCTCTAAACGGGTCTTGAGGGGTTTTTTGCTGAAAGCGAGACTAAGCTT
TAAACTTCGGGTCATAGCTGTTTCCTG

Spytag3-super-TagRFP:

GTAAAACGACGGCCAGTAGCGCTATTAAAGCTTCGAAATTAATACGACTCACTATA
GGGAGACCACAACGGTTTCCCTCTAGAAATAATTTTGTTTAACTTTAAGAAGGAGAT
ATACATATGATGGGACGGGGCGTGCCACATATTGTAATGGTAGACGCATATAAGC
GGTATAAGTCGGGTGGTGGTTCAGGTATGGTTAGTAAGGGCGAAGAACTTATTAA
GGAGAATATGCATATGAAATTGTATATGGAGGGGACTGTTAATAACCACCATTTTAA
ATGCACTTCAGAAGGCGAGGGGAAGCCTTATGAAGGACACAGACCATGCGTATT
AAGGTAGTGGAAGGTGGGCCGCTTCCTTTCGCCTTCGATATTCTGGCAACATCATT
CATGTATGGTTCACGCACTTTCATTAACCACACCCAGGGGATTCCAGACTTTTTAA
ACAGTCGTTCCCGGAAGGCTTCACTTGGGAACGGGTTACTACCTATGAGGATGGC
GGGGTACTCACTGCCACGCAGGACACTTCTCTTCAAGATGGGTGCCTCATCTACA
ATGTGAAAATCCGTGGGGTCAACTTTCCGAGTAATGGTCCGGTTATGCAGAAAAAA
ACTCTTGGTTGGGAAGCCAACACAGAAATGCTCTATCCTGCCGACGGGGGCCTCG
AAGGTCGGACAGTTATGGCTTTGAAGCTGGTCGGTGGTGGGCATCTGATTTGTAA
CTTTAAAACTACTTATCGCTCAAAAAAACCAGCAAAGAACCTGAAAATGCCTGGGG
TCTACTATGTGGATCACCGGCTGGAGCGTATCAAAGAGGCCGACAAAGAAACTTAT
GTAGAGCAACATGAGGTAGCTGTGGCGCGCTATTGTGACTTGCCATCCAAATTGG
GCCACAAGCTCAACGGTATGGATGAACTCTATAAATGGAGCCACCCTCAGTTCGA
GAAATAATAAGTCGACCGGCTGCTAACAAAGCCCGAAAGGAAGCTGAGTTGGCTG
CTGCCACCGCTGAGCAATAACTAGCATAACCCCTTGGGGCCTCTAAACGGGTCTT
GAGGGGTTTTTTGCTGAAAGCGAGACTAAGCTTTAAACTTCGGGTCATAGCTGTTT
CCTG

Spytag3-mKate2:

GTAAAACGACGGCCAGTAGCGCTATTAAAGCTTCGAAATTAATACGACTCACTATA
GGGAGACCACAACGGTTTCCCTCTAGAAATAATTTTGTTTAACTTTAAGAAGGAGAT

ATACATATGGGACGTGGTGTGCCACACATTGTAATGGTTGACGCTTATAAACGCTA
CAAGAGTGGCGGCGGGTCCGGTATGGTATCCGAACTCATCAAAGAGAACATGCAC
ATGAAGTTATACATGGAAGGGACTGTGAATAATCACCACTTTAAGTGTACATCAGA
GGGGGAAGGCAAGCCATATGAGGGTACTCAAACTATGCGCATCAAGGCAGTTGAA
GGGGGTCCTTTACCATTCGCCTTCGATATCTTGGCAACGTCTTTCATGTATGGCAG
TAAGACCTTCATTAACCATACTCAGGGTATTCCTGACTTCTTTAAACAAAGTTTCCC
GGAGGGTTTTACTTGGGAGCGCGTAACTACATATGAGGATGGTGGCGTCCTGACC
GCCACCCAAGACACGTCCCTTCAAGATGGGTGCCTTATCTATAATGTTAAGATCCG
TGGTGTAAATTTTCCGAGCAATGGTCCAGTAATGCAGAAGAAAACTCTGGGTTGGG
AAGCCTCCACTGAAACATTATACCCTGCGGACGGCGGGCTGGAAGGCCGTGCGG
ACATGGCCCTCAAGTTGGTTGGGGGTGGTCATTTAATTTGTAACCTCAAAACTACA
TACCGGTCTAAGAAACCGGCGAAGAATCTGAAAATGCCTGGGGTTTATTACGTGGA
TCGTCGTCTGGAGCGCATTAAGGAAGCGGACAAAGAGACTTACGTGGAGCAACAT
GAGGTGGCCGTCGCACGTTACTGCGATTTGCCTTCTAAGCTGGGTCACCGCTGGT
CCCATCCACAATTCGAGAAATAATAAGTCGACCGGCTGCTAACAAAGCCCGAAAG
GAAGCTGAGTTGGCTGCTGCCACCGCTGAGCAATAACTAGCATAACCCCTTGGGG
CCTCTAAACGGGTCTTGAGGGGTTTTTTGCTGAAAGCGAGACTAAGCTTTAAACTT
CGGGTCATAGCTGTTTCCTG

Spytag3-TagRFP:

GTAAAACGACGGCCAGTAGCGCTATTAAAGCTTCGAAATTAATACGACTCACTATA
GGGAGACCACAACGGTTTCCCTCTAGAAATAATTTTGTTTAACTTTAAGAAGGAGAT
ATACATATGGGACGGGGGGTACCACATATCGTAATGGTGGATGCTTACAAGCGGT
ACAAGAGCGGGGGTGGCAGCGGTATGTCAGAGTTAATCAAAGAGAACATGCACAT
GAAGCTTTACATGGAGGGTACTGTAAACAACCATCACTTTAAGTGCACCAGTGAAG
GGGAGGGTAAACCATATGAGGGTACGCAGACAATGCGCATCAAGGTAGTAGAGG
GCGGCCCGTTACCGTTTGCATTTGACATTCTCGCAACGAGCTTTATGTATGGCTCA
CGGACGTTCATCAATCACACGCAGGGTATCCCAGACTTTTTTAAGCAAAGCTTCCC
GGAGGGCTTTACATGGGAGCGGGTCACAACTTACGAAGATGGTGGGGTGTTGACT
GCAACTCAAGACACCAGCCTCCAGGATGGTTGTCTCATTTATAACGTTAAGATCCG
TGGTGTCAACTTCCCGTCTAATGGGCCAGTGATGCAGAAAAAAACGTTGGGCTGG
GAGGCGAATACGGAGATGTTGTATCCGGCAGACGGTGGCCTTGAAGGTCGGTCA
GACATGGCACTGAAGTTGGTGGGCGGGGGCCACTTAATCTGTAACTTCAAGACGA
CCTATCGCTCGAAAAAACCAGCTAAAAATCTGAAGATGCCAGGTGTTTATTACGTG
GACCACCGGCTGGAGCGTATTAAAGAGGCAGACAAGGAAACCTACGTCGAGCAAC
ACGAGGTTGCCGTAGCCCGCTACTGCGATTTGCCGAGTAAACTTGGTCACAAGTG
GAGCCATCCTCAGTTCGAAAAATAATAAGTCGACCGGCTGCTAACAAAGCCCGAAA
GGAAGCTGAGTTGGCTGCTGCCACCGCTGAGCAATAACTAGCATAACCCCTTGGG
GCCTCTAAACGGGTCTTGAGGGGTTTTTTGCTGAAAGCGAGACTAAGCTTTAAACT
TCGGGTCATAGCTGTTTCCTG

Spytag3-CyOFP1:

GTAAAACGACGGCCAGTAGCGCTATTAAAGCTTCGAAATTAATACGACTCACTATA
GGGAGACCACAACGGTTTCCCTCTAGAAATAATTTTGTTTAACTTTAAGAAGGAGAT
ATACATATGGGACGTGGTGTTCCTCATATTGTAATGGTAGACGCGTATAAACGGTA
CAAGTCAGGCGGGGGTTCGGGGATGGTTTCTAAGGGGGAAGAGTTAATCAAGGAA
AACATGCGCAGCAAATTGTATCTTGAAGGCTCTGTCAATGGGCATCAGTTTAAATG
TACCCATGAGGGTGAAGGGAAACCATATGAAGGTAAACAGACCAATCGTATTAAGG
TCGTCGAGGGGGGGCCGCTCCCATTCGCGTTCGACATTTTAGCAACCCACTTCAT
GTATGGGAGCAAAGTATTCATTAAGTACCCAGCCGATTTACCTGACTACTTTAAGC
AGAGTTTCCCTGAAGGTTTTACATGGAACGCGTAATGGTTTTTGAAGATGGGGGT
GTTCTGACCGCCACGCAAGACACCTCACTGCAAGATGGGGAACTGATCTATAATGT
GAAGGTACGCGGGGTAAACTTCCCGGCAAATGGGCCGGTCATGCAGAAGAAGAC
ACTTGGTTGGGAACCGTCGACAGAGACAATGTACCCTGCAGATGGCGGCCTTGAG
GGTCGCTGCGACAAGGCTTTGAAGCTGGTAGGCGGTGGTCATTTACACGTTAACT
TTAAGACGACATATAAGTCAAAAAAACCTGTCAAGATGCCAGGGGTCCACTATGTT
GATCGTCGGCTGGAACGCATCAAGGAAGCTGATAATGAGACTTATGTAGAACAGTA
CGAACACGCAGTTGCACGTTATTCGAATCTTGGTGGGGGGATGGATGAGCTGTAT
AAGTGGTCTCATCCGCAATTTGAGAAATAATAAGTCGACCGGCTGCTAACAAAGCC
CGAAAGGAAGCTGAGTTGGCTGCTGCCACCGCTGAGCAATAACTAGCATAACCCC
TTGGGGCCTCTAAACGGGTCTTGAGGGGTTTTTTGCTGAAAGCGAGACTAAGCTTT
AAACTTCGGGTCATAGCTGTTTCCTG

Notag-mscarlet:

GTAAAACGACGGCCAGTAGCGCTATTAAAGCTTCGAAATTAATACGACTCACTATA
GGGAGACCACAACGGTTTCCCTCTAGAAATAATTTTGTTTAACTTTAAGAAGGAGAT
ATACATATGGTGTCAAAAGGTGAGGCAGTGATCAAGGAGTTCATGCGTTTCAAAGT
CCACATGGAAGGGTCCATGAATGGCCACGAGTTCGAGATTGAGGGGGAAGGTGA
AGGTCGGCCATATGAGGGTACCCAAACCGCAAAATTGAAGGTCACAAAAGGGGGG
CCACTCCCGTTCTCCTGGGATATTCTGAGCCCTCAGTTCATGTACGGCAGCCGCG
CCTTCACTAAGCACCCTGCCGATATCCCAGACTACTATAAGCAATCATTCCCTGAA
GGGTTTAAATGGGAGCGTGTGATGAACTTCGAGGATGGGGGTGCAGTGACGGTTA
CACAGGATACGTCGTTAGAGGATGGCACCTTGATCTACAAGGTGAAGTTGCGGGG
GACGAACTTTCCACCTGACGGTCCTGTGATGCAGAAAAAAACGATGGGGTGGGAA
GCCTCAACAGAGCGTTTGTACCCTGAGGACGGTGTGTTGAAAGGGGATATCAAGA
TGGCATTACGTCTCAAGGATGGCGGCCGGTACTTGGCCGACTTTAAAACAACCTAT
AAGGCGAAGAAGCCTGTGCAAATGCCTGGGGCTTACAATGTTGATCGCAAACTCG
ACATTACCTCGCACAATGAAGATTACACTGTAGTAGAGCAGTACGAGCGTAGTGAG
GGCCGTCATAGCACGGGTGGTATGGATGAACTTTATAAGTGGAGCCATCCGCAGT
TCGAAAAATAATAAGTCGACCGGCTGCTAACAAAGCCCGAAAGGAAGCTGAGTTG
GCTGCTGCCACCGCTGAGCAATAACTAGCATAACCCCTTGGGGCCTCTAAACGGG

TCTTGAGGGGTTTTTTGCTGAAAGCGAGACTAAGCTTTAAACTTCGGGTCATAGCT
GTTTCCTG

mScarlet3-spytag003:

GTAAAACGACGGCCAGTAGCGCTATTAAAGCTTCGAAATTAATACGACTCACTATA
GGGAGACCACAACGGTTTCCCTCTAGAAATAATTTTGTTTAACTTTAAGAAGGAGAT
ATACATATGGGTCGTGGTGTACCTCATATCGTAATGGTGGATGCGTATAAGTCCGG
CGGGGGGTCTGGGGATAGCACGGAGGCTGTAATCAAGGAGTTCATGCGTTTCAAA
GTACATATGGAAGGGTCTATGAACGGGCATGAATTTGAGATCGAAGGGGAGGGCG
AAGGTCGGCCATACGAGGGCACACAAACAGCGAAACTCCGGGTTACAAAAGGTGG
GCCTCTTCCTTTCTCGTGGGACATCCTTAGTCCTCAATTCATGTATGGCTCGCGTG
CCTTTACAAAGCACCCTGCCGACATCCCTGATTACTGGAAGCAAAGCTTCCCGGAA
GGGTTCAAATGGGAGCGCGTCATGAATTTTGAAGATGGTGGCGCCGTATCAGTGG
CTCAAGATACGTCTCTTGAAGACGGCACACTGATCTACAAGGTGAAATTACGGGGC
ACGAATTTCCCACCAGATGGTCCAGTCATGCAGAAAAAGACAATGGGGTGGGAAG
CCTCAACCGAGCGCCTGTACCCGGAGGATGTCGTGTTAAAGGGTGATATCAAGAT
GGCACTTCGCTTGAAAGACGGGGGTCGTTATCTCGCAGACTTCAAAACAACCTATC
GTGCTAAAAAGCCGGTTCAAATGCCAGGCGCATTCAATATTGATCGTAAGCTGGAT
ATCACATCGCATAACGAAGATTATACAGTAGTGGAACAATATGAACGGTCAGTCGC
TCGGCACAGTACGGGCGGTTCAGGGGGGAGCTGGAGCCATCCGCAGTTCGAAAA
ATAATAAGTCGACCGGCTGCTAACAAAGCCCGAAAGGAAGCTGAGTTGGCTGCTG
CCACCGCTGAGCAATAACTAGCATAACCCCTTGGGGCCTCTAAACGGGTCTTGAG
GGGTTTTTTGCTGAAAGCGAGACTAAGCTTTAAACTTCGGGTCATAGCTGTTTCCT
G

Nb6101-mScarlet3:

GTAAAACGACGGCCAGTAGCGCTATTAAAGCTTCGAAATTAATACGACTCACTATA
GGGAGACCACAACGGTTTCCCTCTAGAAATAATTTTGTTTAACTTTAAGAAGGAGAT
ATACAT ATG CGC GTC CAA TTA GTT GAG AGT GGT GGC GGC TTA GTA CAG
GCG GGC GGG TCT CTC CGT CTG AGC TGT GCG GCC GAT GGC TTT GAT TTC
AAA AGC TAT GCT ATG GGT TGG TAT CGG CAG GCT CCG GGT CGC GAA GAT
GAA CTC GTC GCG GCA ATC ACG GCA TCG GGT GAT TAT ACC TAT TAT GCG
GAT TCT GTA AAG GGC CGC TTC ACG ATT TCG CGG GAT AAC GCT AAA AAC
ACT GTT TAC TTG CAA ATG AAT AGT TTA AAA CCG GAT GAT ACG GCG GTG
TAT TAT TGT GCT GCG CTG TCC TAT GTG GCG GAG GGT TAT TGG GGT CAG
GGG ACG CAA GTC ACT GTT AGC TCG AGT GGC GGT GGT AGT GGT GAC TCA
ACG GAA GCG GTA ATC AAA GAA TTT ATG CGC TTC AAA GTC CAT ATG GAG
GGC AGC ATG AAC GGT CAT GAA TTT GAG ATC GAG GGC GAA GGT GAG GGC
CGC CCG TAT GAG GGG ACG CAG ACG GCG AAA CTC CGT GTC ACA AAA GGC
GGG CCA TTG CCG TTT TCC TGG GAT ATT CTG TCG CCG CAG TTT ATG TAT

GGC TCC CGC GCG TTC ACC AAA CAC CCA GCC GAT ATC CCG GAT TAC TGG AAA CAG TCC TTC CCG GAG GGT TTT AAA TGG GAG CGC GTA ATG AAC TTT GAA GAT GGT GGC GCC GTG TCA GTT GCT CAG GAC ACC AGC TTG GAG GAC GGT ACC CTT ATT TAT AAA GTG AAA CTG CGC GGG ACC AAT TTT CCA CCA GAT GGC CCG GTC ATG CAG AAA AAA ACC ATG GGC TGG GAA GCA TCT ACC GAG CGT CTG TAC CCA GAG GAC GTG GTC CTG AAA GGT GAT ATC AAA ATG GCC CTC CGC CTG AAG GAC GGC GGT CGC TAT CTG GCG GAT TTT AAG ACG ACC TAT CGC GCC AAG AAA CCG GTT CAA ATG CCG GGC GCT TTC AAT ATT GAT CGT AAA TTG GAT ATT ACA TCT CAC AAT GAA GAT TAT ACC GTT GTG GAA CAA TAT GAA CGC AGT GTC GCG CGG CAT AGC ACA GGG GGC TCC GGC GGC TCC
TGGAGCCATCCGCAGTTCGAAAAATAATAAGTCGACCGGCTGCTAACAAAGCCCG AAAGGAAGCTGAGTTGGCTGCTGCCACCGCTGAGCAATAACTAGCATAACCCCTT GGGGCCTCTAAACGGGTCTTGAGGGGTTTTTTGCTGAAAGCGAGACTAAGCTTTAA ACTTCGGGTCATAGCTGTTTCCTG

NB457 -mScarlet3

GTAAAACGACGGCCAGTAGCGCTATTAAAGCTTCGAAATTAATACGACTCACTATA GGGAGACCACAACGGTTTCCCTCTAGAAATAATTTTGTTTAACTTTAAGAAGGAGAT ATACAT ATG GGC GAA GTT CAG TTG GTG GAA AGC GGC GGC GGC CTG GTT CAG GCA GGC GGG AGT CTT CGC CTG TCT TGT GCG GCG AGC GGC CGT ACG ATT AGT TCG GTC GCT ATG GGT TGG TTC CGC CAG GCA CCG GAG AAG GAA CGT GAA TTC GTG GCC GCG ATT ACT TGG AGC GGG GAC TAT ACC AAC GTG GCC GAT TCT ATG AAG GGT CGT TTC ACC ATC AGC CGT GAT AAC GCT CGG AAA ACC GTT AGC CTG CAG TTG ACG AAT CTC AAA CCT GAA GAC ACC GCT GTG TAC TAT TGT GCA GCG GAT CTG CGC GGT GGC TCC ATT TAT GGT ACA GCA GAT TAT GTC TAC TGG GGT CAG GGC ACC CAG GTC ACA GTG TCC AGC CTC GAG GGC GGT GGG GGC AGC GGC GGC GGT GGC AGT GAC AGT ACC GAA GCG GTC ATC AAA GAA TTT ATG CGT TTT AAA GTT CAT ATG GAA GGG TCT ATG AAT GGT CAC GAG TTT GAG ATT GAA GGT GAA GGC GAA GGT CGG CCA TAT GAA GGC ACC CAG ACG GCG AAA CTG CGC GTG ACC AAA GGC GGT CCG CTG CCT TTC AGT TGG GAT ATT TTA AGC CCG CAG TTT ATG TAC GGC AGC CGG GCA TTC ACC AAA CAT CCG GCC GAC ATC CCG GAT TAC TGG AAG CAA TCA TTT CCG GAA GGT TTT AAG TGG GAA CGT GTG ATG AAC TTT GAG GAT GGG GGC GCC GTT AGT GTG GCG CAA GAT ACC TCC CTG GAG GAT GGT ACC CTG ATT TAT AAA GTG AAA TTA CGC GGT ACC AAT TTC CCG CCG GAC GGG CCG GTA ATG CAG AAA AAG ACC ATG GGC TGG GAG GCA TCT ACC GAG CGG CTG TAT CCG GAA GAT GTA GTA TTG AAA GGT GAT ATC AAA ATG GCC CTG CGC CTC AAA GAC GGG GGT CGT TAT TTA GCG GAT TTT AAA ACC ACC TAT CGT GCC AAA AAG CCA GTG CAG ATG CCA GGG GCG TTT AAC ATT GAT

CGG AAA TTA GAC ATC ACT TCA CAT AAT GAA GAC TAC ACG GTT GTG GAA CAG TAT GAA CGC AGC GTG GCC CGT CAC TCC ACC GGC GGT AGT GGC GGT AGTTGGAGCCATCCGCAGTTCGAAAAATAATAAGTCGACCGGCTGCTAACAAAGC CCGAAAGGAAGCTGAGTTGGCTGCTGCCACCGCTGAGCAATAACTAGCATAACCC CTTGGGGCCTCTAAACGGGTCTTGAGGGGTTTTTTGCTGAAAGCGAGACTAAGCTT TAAACTTCGGGTCATAGCTGTTTCCTG

ZF456-linker-mCitrine

GTAAAACGACGGCCAGTAGCGCTATTAAAGCTTCGAAATTAATACGACTCACTATA GGGAGACCACAACGGTTTCCCTCTAGAAATAATTTTGTTTAACTTTAAGAAGGAGAT ATACAT ATG GAG GGG CGT CGT AGC GAC ACC TGC GAA TAT TGC GGT AAA GTA TTC AAA AAC TGT TCT AAC CTG ACA GTG CAC CGC CGC TCG CAT ACG GGC GAA CGG CCG TAT AAA TGT GAG CTG TGT AAT TAC GCT TGC GCG CAA TCC TCA AAA TTG ACA CGC CAT ATG AAA ACC CAC GGT CAA GTG GGT AAG GAT GTG TAC AAA TGC GAA ATT TGC AAA ATG CCG TTT TCT GTG TAT TCG ACT TTG GAA AAA CAC ATG AAG AAG TGG CAC AGT GAC CGC GTA CTT AAC AAC GAT ATT AAA ACC GAA GGT GGG GGC GGT AGC GGT GGC GGG GGG TCT GTG AGC AAA GGT GAG GAG CTG TTC ACG GGC GTA GTT CCA ATT CTC GTT GAA TTG GAT GGC GAT GTG AAC GGC CAT AAA TTT AGC GTC TCC GGG GAG GGT GAG GGG GAT GCT ACC TAT GGG AAA CTT ACG CTG AAG TTT ATT TGT ACT ACC GGC AAA CTT CCA GTT CCG TGG CCT ACC CTG GTG ACC ACT TTC GGG TAT GGC TTG ATG TGT TTT GCA CGC TAC CCG GAT CAC ATG AAA CAA CAC GAT TTT TTC AAG AGC GCA ATG CCT GAA GGT TAT GTT CAG GAA CGC ACC ATT TTT TTC AAA GAC GAT GGT AAT TAT AAG ACG CGC GCA GAA GTG AAA TTT GAA GGC GAC ACC CTG GTA AAT CGC ATC GAA TTA AAA GGT ATT GAC TTC AAA GAA GAT GGT AAT ATC TTG GGT CAT AAA CTG GAA TAT AAT TAT AAC AGC CAT AAC GTT TAT ATT ATG GCG GAT AAA CAG AAG AAC GGT ATC AAA GTT AAC TTC AAG ATC CGT CAT AAT ATC GAA GAC GGC TCC GTT CAG CTG GCC GAT CAT TAC CAG CAG AAT ACA CCG ATT GGG GAC GGT CCA GTG TTG CTG CCA GAT AAT CAC TAT CTG TCT TAT CAG AGT AAA CTC TCC AAA GAT CCA AAT GAA AAA CGG GAT CAT ATG GTC TTG TTA GAG TTT GTT ACA GCA GCG GGC ATT ACC CTC GGT ATG GAT GAA CTG TAC AAATGGAGCCATCCGCAGTTCGAAAAATAATAAGTCGACCGGCTGCTAACAAAGCC CGAAAGGAAGCTGAGTTGGCTGCTGCCACCGCTGAGCAATAACTAGCATAACCCC TTGGGGCCTCTAAACGGGTCTTGAGGGGTTTTTTGCTGAAAGCGAGACTAAGCTTT AAACTTCGGGTCATAGCTGTTTCCTG

ZF456-linker-mTurquoise2

GTAAAACGACGGCCAGTAGCGCTATTAAAGCTTCGAAATTAATACGACTCACTATA GGGAGACCACAACGGTTTCCCTCTAGAAATAATTTTGTTTAACTTTAAGAAGGAGAT

ATACAT ATG GAA GGC CGT CGT AGT GAT ACG TGT GAA TAC TGC GGT AAA GTA TTT AAA AAT TGC AGT AAC CTT ACC GTG CAT CGT CGC TCC CAC ACG GGG GAA CGG CCG TAC AAA TGT GAA CTT TGT AAT TAT GCT TGC GCA CAA AGT AGT AAA TTG ACG CGT CAT ATG AAA ACG CAC GGG CAG GTG GGT AAA GAT GTG TAT AAA TGC GAA ATT TGC AAA ATG CCG TTT AGC GTT TAT AGC ACG CTG GAG AAA CAT ATG AAG AAA TGG CAC TCT GAT CGT GTC CTG AAT AAT GAC ATC AAA ACT GAA GGC GGT GGG GGT TCG GGT GGG GGC GGC TCT GTG TCG AAA GGC GAA GAG CTG TTT ACC GGC GTG GTG CCG ATC CTC GTT GAA CTT GAT GGC GAT GTT AAT GGT CAT AAA TTT TCG GTC AGT GGC GAG GGC GAG GGT GAT GCG ACC TAT GGC AAA TTG ACC CTG AAG TTC ATC TGT ACC ACT GGT AAA CTG CCA GTG CCA TGG CCG ACC CTC GTC ACA ACA TTA AGC TGG GGC GTC CAG TGC TTC GCG CGG TAT CCA GAC CAT ATG AAA CAG CAC GAT TTT TTT AAA TCC GCG ATG CCA GAA GGT TAC GTT CAG GAG CGT ACC ATC TTC TTC AAA GAT GAT GGT AAT TAC AAA ACC CGT GCC GAG GTA AAA TTC GAG GGT GAC ACC CTG GTG AAC CGC ATC GAA CTG AAA GGC ATT GAC TTT AAA GAA GAC GGT AAC ATT TTG GGT CAT AAG CTC GAA TAC AAC TAC TTT AGT GAC AAC GTC TAC ATC ACG GCG GAT AAA CAG AAA AAT GGT ATT AAG GCA AAT TTC AAA ATT CGC CAC AAT ATT GAG GAC GGC GGC GTG CAG TTA GCG GAT CAC TAC CAG CAG AAT ACG CCA ATT GGG GAC GGC CCG GTA TTA CTT CCG GAT AAC CAC TAT CTG AGC ACT CAG AGC AAA CTG AGC AAA GAT CCT AAC GAG AAA CGC GAT CAC ATG GTG CTG CTC GAG TTT GTT ACA GCC GCT GGG ATT ACG TTA GGT ATG GAT GAG TTG TAT AAATGGAGCCATCCGCAGTTCGAAAAATAATAAGTCGACCGGCTGCTAACAAAGCCCGAAAGGAAGCTGAGTTGGCTGCTGCCACCGCTGAGCAATAACTAGCATAACCCCTTGGGGCCTCTAAACGGGTCTTGAGGGGTTTTTTGCTGAAAGCGAGACTAAGCTTTAAACTTCGGGTCATAGCTGTTTCCTG

sdAb-1E2-mScarlet3:

GTAAAACGACGGCCAGTAGCGCTATTAAAGCTTCGAAATTAATACGACTCACTATAGGGAGACCACAACGGTTTCCCTCTAGAAATAATTTTGTTTAACTTTAAGAAGGAGATATACAT ATG CAG GTG CAG CTG GTT GAG AGT GGC GGC GGT CTG GTT CAA CCT GGT GGC TCT CTG CGC TTG TCA TGC GCA GCA TCC GGC GCG GAT GCC TTT GTA ATT ATC GGC GGG TGG TTT CGT CAA GCG CCG GGT AAA GGT CTG GAA GCG GTG GCT GCG ATT GCG TGG ACG GAT CAG CAT GAA TAT TAC GCA GAT TCG GTA AAG GGT CGT TTT ACC ATC TCG CGC GAC AAT AGC AAA AAC ACG CTG TAC CTG CAA ATG AAC TCT TTA CGC GCG GAA GAT ACC GCG GTC TAT TAC TGC GCT GCG CAG GAT AGT GCA TAC ATT AAG TCT AAA GGT AGT CGC GCA TAT GAA TAT TGG GGT CAG GGC ACC CAG GTT ACC GTC TCC AGC GGT GGG GGT GGG AGC GGG GGC GGT GGG AGC GAC AGT ACG GAG GCG GTA ATT AAG GAA TTT ATG CGT TTT AAG GTG CAT ATG GAA GGC TCG ATG

AAT GGC CAC GAA TTT GAG ATC GAA GGC GAA GGT GAA GGC CGT CCT TAC GAA GGT ACC CAG ACC GCG AAA CTG CGC GTA ACT AAA GGG GGT CCG CTC CCG TTC AGC TGG GAT ATC CTG TCG CCG CAA TTC ATG TAT GGT AGC CGC GCG TTT ACT AAA CAT CCT GCA GAT ATT CCA GAT TAC TGG AAA CAG AGC TTC CCG GAA GGC TTT AAA TGG GAA CGG GTG ATG AAT TTC GAA GAT GGT GGT GCA GTC AGC GTC GCG CAA GAT ACG TCG CTG GAA GAT GGC ACT CTC ATT TAT AAG GTG AAA CTT CGC GGC ACC AAT TTC CCA CCA GAT GGC CCG GTG ATG CAA AAA AAA ACT ATG GGC TGG GAA GCA AGT ACC GAA CGT CTG TAT CCA GAA GAT GTT GTT CTG AAA GGT GAT ATC AAA ATG GCG CTG CGC CTG AAA GAT GGC GGT CGT TAT TTA GCG GAT TTC AAA ACC ACG TAT CGC GCA AAA AAA CCG GTT CAA ATG CCA GGG GCG TTC AAC ATT GAT CGG AAA CTG GAT ATC ACA TCA CAT AAC GAG GAT TAC ACC GTC GTC GAA CAG TAC GAA CGT AGT GTC GCA CGC CAC TCG ACG GGG GGT TCG GGT GGG AGT TGGAGCCATCCGCAGTTCGAAAAATAATAAGTCGACCGGCTGCTAACAAAGCCCG AAAGGAAGCTGAGTTGGCTGCTGCCACCGCTGAGCAATAACTAGCATAACCCCTT GGGGCCTCTAAACGGGTCTTGAGGGGTTTTTTGCTGAAAGCGAGACTAAGCTTTAA ACTTCGGGTCATAGCTGTTTCCTG

sdAb-2F2-mScarlet3:

GTAAAACGACGGCCAGTAGCGCTATTAAAGCTTCGAAATTAATACGACTCACTATA GGGAGACCACAACGGTTTCCCTCTAGAAATAATTTTGTTTAACTTTAAGAAGGAGAT ATACAT ATG CAG GTT CAG CTC GTT GAG TCT GGT GGC GGT TTA GTA CAG CCG GGC GGC TCG TTG CGC CTG AGC TGC GCG GCA AGC GGG TTG GCT CAG TCG AAG TGG GCA TAT GGC TGG TTT CGT CAG GCG CCG GGC AAA GGC TTG GAA GCG GTG GCG GCA ATT GAT GTA GCC ACA GGG CCG TGG TAT TAC GCG GAC AGC GTA AAA GGC CGC TTT ACC ATT TCA CGT GAT AAT TCT AAG AAT ACA CTG TAC TTA CAG ATG AAC AGC TTG CGT GCG GAA GAT ACG GCC GTG TAT TAC TGT GCG GCG CAC CAT ATT CCA ACA AAG CAC CCG GCA TTC CCA GAT TTT CGT GAT TAC TGG GGT CAA GGC ACG CAA GTC ACC GTT TCG TCT GGC GGT GGG GGG AGC GGG GGT GGT GGG TCG GAT TCC ACG GAA GCC GTC ATT AAA GAG TTC ATG CGT TTT AAA GTA CAT ATG GAA GGC AGT ATG AAC GGC CAC GAA TTC GAA ATC GAG GGT GAA GGC GAG GGT CGT CCT TAT GAA GGT ACT CAG ACC GCC AAA CTT CGC GTT ACG AAG GGG GGT CCG TTA CCG TTT TCC TGG GAT ATT TTA TCG CCG CAA TTT ATG TAC GGG AGC CGC GCA TTT ACC AAA CAT CCT GCG GAC ATT CCA GAT TAC TGG AAG CAA AGC TTT CCG GAA GGT TTT AAG TGG GAG CGT GTC ATG AAT TTC GAA GAT GGG GGC GCA GTA AGC GTG GCG CAG GAT ACG AGT CTG GAA GAT GGT ACG CTG ATC TAT AAA GTC AAA TTA CGC GGC ACA AAT TTT CCT CCT GAT GGT CCA GTT ATG CAA AAG AAG ACC ATG GGC TGG GAG GCC TCC ACT GAG CGC CTG TAT CCG GAA GAT GTG GTA CTT AAG GGG GAT ATT AAA ATG GCA TTA CGC

CTG AAA GAC GGC GGC CGC TAT CTG GCC GAT TTT AAG ACG ACC TAT CGC GCG AAA AAG CCG GTT CAA ATG CCA GGC GCT TTT AAC ATC GAT CGC AAA CTG GAT ATC ACC AGC CAC AAT GAG GAC TAT ACC GTC GTT GAA CAG TAT GAG CGC AGT GTG GCT CGC CAC AGC ACC GGC GGC AGC GGT GGT AGC

TGGAGCCATCCGCAGTTCGAAAAATAATAAGTCGACCGGCTGCTAACAAAGCCCG AAAGGAAGCTGAGTTGGCTGCTGCCACCGCTGAGCAATAACTAGCATAACCCCTT GGGGCCTCTAAACGGGTCTTGAGGGGTTTTTTGCTGAAAGCGAGACTAAGCTTTAA ACTTCGGGTCATAGCTGTTTCCTG

sdAb-5F8-mScarlet3:

GTAAAACGACGGCCAGTAGCGCTATTAAAGCTTCGAAATTAATACGACTCACTATA GGGAGACCACAACGGTTTCCCTCTAGAAATAATTTTGTTTAACTTTAAGAAGGAGAT ATACAT ATG CAG GTT CAA TTG GTG GAG AGC GGT GGG GGC CTT GTG CAG CCG GGT GGG TCA CTG CGT TTG AGC TGC GCG GCG TCC GGT CGT CAG GCT GTA GTG TTT GGC TGG TTT CGT CAG GCG CCG GGC AAA GGT CTG GAA GCG GTG GCA GCC ATT CAT GTC CCG GCA AAA AAC CGG TAT TAT GCG GAT AGT GTA AAA GGG CGG TTT ACG ATT TCC CGC GAT AAT TCC AAA AAT ACG CTT TAT CTC CAG ATG AAC TCC CTC CGC GCG GAA GAT ACA GCG GTG TAT TAC TGT GCA GCA CAT TAT GAA TTC AAT GAC TTC GTC TGG CAG GGC TAT AGT AGC GAC TAT TGG GGG CAG GGG ACC CAG GTT ACG GTT TCG AGC GGT GGG GGT GGG TCA GGT GGG GGC GGT AGC GAC AGC ACC GAG GCT GTA ATT AAG GAA TTC ATG CGC TTT AAG GTA CAT ATG GAG GGT TCC ATG AAT GGT CAT GAG TTT GAA ATT GAG GGG GAG GGC GAA GGT CGC CCT TAT GAA GGC ACT CAG ACG GCG AAA TTG CGG GTC ACG AAG GGC GGG CCA TTG CCG TTT AGT TGG GAT ATC TTA TCT CCG CAG TTT ATG TAT GGC AGC CGC GCC TTC ACG AAA CAT CCG GCC GAT ATT CCG GAC TAC TGG AAA CAA AGT TTT CCG GAA GGT TTC AAA TGG GAG CGC GTA ATG AAC TTT GAA GAC GGT GGC GCA GTT TCC GTG GCA CAG GAC ACC TCG TTG GAA GAC GGG ACG TTG ATT TAT AAA GTT AAG TTG CGC GGT ACC AAC TTT CCG CCT GAT GGC CCG GTT ATG CAG AAA AAA ACC ATG GGT TGG GAA GCC TCT ACG GAA CGG CTG TAT CCG GAG GAT GTG GTC CTC AAA GGC GAT ATT AAA ATG GCG TTA CGC CTG AAG GAC GGC GGT CGT TAT CTG GCG GAT TTT AAG ACG ACA TAT CGC GCA AAG AAG CCG GTC CAG ATG CCA GGC GCG TTT AAT ATT GAC CGT AAG TTA GAT ATT ACC TCA CAC AAT GAA GAC TAT ACA GTT GTA GAA CAG TAC GAA CGT TCG GTT GCC CGT CAC AGC ACG GGC GGT TCA GGC GGT AGT

TGGAGCCATCCGCAGTTCGAAAAATAATAAGTCGACCGGCTGCTAACAAAGCCCG AAAGGAAGCTGAGTTGGCTGCTGCCACCGCTGAGCAATAACTAGCATAACCCCTT

GGGGCCTCTAAACGGGTCTTGAGGGGTTTTTTGCTGAAAGCGAGACTAAGCTTTAA
ACTTCGGGTCATAGCTGTTTCCTG

7R6X_3-mcitrine

GTAAAACGACGGCCAGTAGCGCTATTAAAGCTTCGAAATTAATACGACTCACTATA
GGGAGACCACAACGGTTTCCCTCTAGAAATAATTTTGTTTAACTTTAAGAAGGAGAT
ATACAT ATG GAA ACT GGC ACA CGC TTC CCA AAC ATC ACC AAT CTT TGT
CCG TTT GGT GAA GTG TTC AAT GCG ACT CGT TTT GCG AGC GTG TAT GCG
TGG AAT CGG AAA CGC ATT AGC AAT TGC GTG GCC GAT TAT AGT GTA TTA
TAT AAC AGT GCT TCA TTT TCG ACC TTT AAA TGC TAC GGC GTC TCT CCA
ACG AAA CTT AAC GAT TTG TGC TTC ACC AAC GTT TAC GCA GAT TCC TTC
GTG ATT CGC GGG GAT GAG GTA CGT CAA ATC GCG CCA GGC CAG ACT GGC
AAG ATT GCC GAT TAT AAC TAC AAA CTC CCG GAT GAC TTT ACC GGC TGC
GTT ATC GCT TGG AAT TCA AAT AAC CTG GAT TCA AAA GTG GGC GGT AAC
TAT AAC TAC CTG TAT CGC TTA TTT CGT AAG AGT AAC CTG AAA CCG TTT GAA
CGC GAT ATT TCC ACA GAA ATT TAC CAG GCA GGC TCC ACC CCA TGC AAT
GGC GTC GAA GGT TTT AAT TGT TAT TTC CCA CTT CAA AGC TAC GGC TTC
CAA CCA ACA AAT GGT GTA GGT TAC CAA CCG TAC CGC GTG GTG GTG CTG
TCG TTT GAG CTG TTA CAT GCT CCG GCT ACC GTG TGC GGC CCA AAA AAA
AGT ACC CAC CAC CAT CAT CAT CAT CAC CAC GGT GGT GGT GGG AGT GGC
GGT GGT GGT TCC GTG TCG AAA GGT GAA GAG CTG TTC ACG GGC GTT GTC
CCG ATC TTA GTG GAA CTC GAT GGC GAC GTT AAT GGC CAT AAG TTT TCT
GTC TCT GGC GAG GGG GAA GGT GAT GCC ACC TAT GGG AAA TTA ACC CTT
AAA TTT ATC TGT ACC ACT GGC AAA TTG CCG GTC CCG TGG CCG ACC CTG
GTG ACA ACC TTT GGT TAT GGT CTG ATG TGC TTT GCC CGC TAT CCG GAC
CAT ATG AAA CAG CAT GAT TTT TTC AAA TCT GCG ATG CCG GAA GGC TAC
GTC CAG GAA CGT ACA ATT TTC TTC AAA GAT GAC GGC AAC TAT AAG ACC
CGG GCC GAA GTG AAG TTC GAG GGT GAT ACA CTT GTA AAC CGT ATC GAA
CTG AAA GGC ATT GAC TTT AAA GAG GAT GGG AAC ATT CTG GGG CAC AAA
CTG GAA TAC AAC TAT AAC TCA CAC AAC GTC TAT ATT ATG GCA GAC AAA
CAG AAG AAT GGG ATC AAA GTG AAT TTC AAA ATT CGC CAT AAT ATT GAA
GAT GGT AGT GTA CAG CTG GCC GAC CAC TAT CAG CAG AAT ACC CCG ATT
GGC GAT GGT CCG GTT CTG TTG CCA GAT AAC CAC TAT CTG TCT TAC CAA
TCA AAA CTG TCA AAG GAT CCG AAT GAA AAA CGC GAC CAC ATG GTA CTG
CTG GAA TTT GTT ACA GCT GCG GGT ATC ACT TTA GGC ATG GAT GAA TTA
TAT AAA

TGGAGCCATCCGCAGTTCGAAAAATAATAAGTCGACCGGCTGCTAACAAAGCCCG
AAAGGAAGCTGAGTTGGCTGCTGCCACCGCTGAGCAATAACTAGCATAACCCCTT

GGGGCCTCTAAACGGGTCTTGAGGGGTTTTTTGCTGAAAGCGAGACTAAGCTTTAA
ACTTCGGGTCATAGCTGTTTCCTG

Spycatcher3-mcitrine:

GTAAAACGACGGCCAGTAGCGCTATTAAAGCTTCGAAATTAATACGACTCACTATA
GGGAGACCACAACGGTTTCCCTCTAGAAATAATTTTGTTTAACTTTAAGAAGGAGAT
ATACAT ATG GTG ACT ACG TTA AGT GGC CTG TCA GGC GAA CAG GGG CCG
TCG GGC GAC ATG ACT ACC GAA GAG GAT TCC GCC ACG CAC ATC AAA TTT
TCT AAA CGC GAC GAA GAT GGT CGT GAA CTC GCC GGT GCA ACC ATG GAA
TTG CGC GAT AGC TCC GGT AAA ACG ATT TCA ACC TGG ATC TCG GAT GGG
CAT GTG AAA GAT TTT TAT TTG TAT CCG GGT AAA TAC ACC TTT GTC GAA ACC
GCG GCA CCG GAT GGT TAT GAG GTG GCA ACG CCG ATT GAA TTT ACG GTC
AAT GAG GAT GGT CAA GTC ACG GTC GAT GGC GAA GCA ACC GAG GGC GAT
GCA CAC ACG GGC GGT GGG GGG TCC GGT GGC GGT GGC TCT ATG GTG TCA
AAA GGT GAG GAG CTC TTC ACC GGC GTT GTG CCG ATT CTG GTA GAA CTC
GAT GGT GAC GTC AAC GGC CAT AAA TTT AGT GTA TCT GGC GAA GGC GAA
GGC GAT GCG ACC TAT GGT AAA CTG ACG TTA AAA TTC ATC TGT ACT ACC
GGG AAA CTG CCG GTT CCG TGG CCA ACC TTA GTG ACT ACG TTC GGC TAT
GGT CTG ATG TGC TTC GCA CGC TAC CCG GAT CAC ATG AAG CAG CAC GAT
TTT TTT AAA TCA GCG ATG CCG GAA GGG TAC GTC CAA GAA CGG ACG ATC
TTT TTC AAA GAC GAT GGC AAT TAC AAG ACG CGC GCA GAA GTA AAA TTT
GAG GGT GAT ACC CTG GTG AAC CGT ATC GAA CTT AAA GGC ATT GAT TTC
AAA GAA GAC GGC AAC ATC CTG GGC CAC AAA CTG GAA TAT AAT TAC AAT
AGT CAT AAC GTT TAT ATT ATG GCC GAC AAA CAG AAA AAT GGT ATC AAA
GTC AAT TTT AAA ATT CGC CAC AAC ATT GAA GAT GGC AGT GTG CAG TTG
GCC GAC CAC TAC CAG CAA AAT ACG CCG ATC GGG GAC GGC CCG GTG CTG
TTA CCG GAC AAC CAC TAT CTG TCC TAC CAG TCA AAG CTG TCT AAA GAC
CCG AAT GAA AAA CGC GAC CAC ATG GTC CTT CTG GAA TTT GTG ACG GCG
GCA GGC ATC ACC CTG GGC ATG GAT GAA CTG TAT AAA

TGGAGCCATCCGCAGTTCGAAAAATAATAAGTCGACCGGCTGCTAACAAAGCCCG
AAAGGAAGCTGAGTTGGCTGCTGCCACCGCTGAGCAATAACTAGCATAACCCCTT
GGGGCCTCTAAACGGGTCTTGAGGGGTTTTTTGCTGAAAGCGAGACTAAGCTTTAA
ACTTCGGGTCATAGCTGTTTCCTG

ZF456 expression Genes:

Codon optimized ZF456-strepII:

5’-
ATGGAAGGCCGCCGCAGCGATACCTGCGAATATTGCGGCAAAGTGTTTAAAAACT
GCAGCAACCTGACCGTGCATCGCCGCAGCCATACCGGCGAACGCCCGTATAAATG
CGAACTGTGCAACTATGCGTGCGCGCAGAGCAGCAAACTGACCCGCCATATGAAA
ACCCATGGCCAGGTGGGCAAAGATGTGTATAAATGCGAAATTTGCAAAATGCCGTT
TAGCGTGTATAGCACCCTGGAAAAACATATGAAAAAATGGCATAGCGATCGCGTGC
TGAACAACGATATTAAAACCGAATGGAGCCATCCGCAGTTTGAAAAATAATAA -3’

FW: (ZF456GA_FW):

CCGCAAGCTTGTCGACGGAGCTCGAATTCGATGGAAGGCCGC

RV:(ZF456GA_RV):

TGGCCATGGATATCGGAATTAATTCGGATCTTATTATTTTTCAAACTGCGG

FW: (ZF456_FW): ATGGAAGGCCGCCGC

RV: (ZF456_RV):ttattaTTTTTCAAACTGCGG

### Section 6: Python scripts used to fit the data to the binding model

#### Sample donor quenching fitting:

```
import numpy as np

import matplotlib.pyplot as plt

from scipy.optimize import curve_fit

import matplotlib.cm as cm

# === Your datasets (raw donor fluorescence) ===

X0 = 0.1   # max acceptor (µM)

D0 = 0.05 # highest donor concentration (µM)

datasets = [

  {"D": D0,

   "A": np.array([X0,X0/2,X0/4,X0/8,X0/16,X0/32,X0/64,0]),

   "F": np.array([13855,16768,24924,27481,30579,29488,32163,30674,])},
```

```
  {"D": D0/2,
   "A": np.array([X0,X0/2,X0/4,X0/8,X0/16,X0/32,X0/64,0]),
   "F": np.array([6474,6951,10026,13122,15588,14971,16418,15786])},
  {"D": D0/4,
   "A": np.array([X0,X0/2,X0/4,X0/8,X0/16,X0/32,X0/64,0]),
   "F": np.array([3685,3578,4163,5617,6475,7581,7894,8052])},
   {"D": D0/8,
   "A": np.array([X0,X0/2,X0/4,X0/8,X0/16,X0/32,X0/64,0]),
   "F": np.array([1660,1767,1659,2346,2882,3376,3720,3877])},
]
# === Binding quenching model ===
def quenching_model(A, Kd, D, Fmax, Fmin):
  """Simple 1:1 donor–acceptor quenching model."""
  frac_bound = 0.5 * ((A + D + Kd) - np.sqrt((A + D + Kd)**2 - 4*A*D)) / D
  return Fmax - (Fmax - Fmin) * frac_bound
# === Global model (shared Kd, individual Fmax/Fmin per dataset) ===
def global_model(X_all, Kd, *params):
  """Concatenated model for all datasets."""
  model = []
  for i, dataset in enumerate(datasets):
    A = dataset["A"]
    D = dataset["D"]
    Fmax, Fmin = params[2*i], params[2*i+1]
    model.extend(quenching_model(A, Kd, D, Fmax, Fmin))
  return np.array(model)

# === Prepare concatenated data ===
Y_all = np.concatenate([d["F"] for d in datasets])
X_all = np.arange(len(Y_all))  # dummy X for fitting

# === Initial guesses ===
```

```
p0 = [0.01]  # initial Kd (µM)
for d in datasets:
    p0 += [np.max(d["F"]), np.min(d["F"])]  # Fmax, Fmin per dataset

# === Bounds ===
lower = [1e-9] + [0, 0]*len(datasets)
upper = [10.0] + [np.inf, np.inf]*len(datasets)

# === Fit ===
popt, _ = curve_fit(global_model, X_all, Y_all, p0=p0, bounds=(lower, upper))
Kd_fit = popt[0]
params_fit = popt[1:]

# === Calculate fit quality ===
Y_fit_all = global_model(X_all, *popt)
SS_res = np.sum((Y_all - Y_fit_all)**2)
SS_tot = np.sum((Y_all - np.mean(Y_all))**2)
R2 = 1 - SS_res/SS_tot

# === Plot ===
plt.style.use('default')
fig, ax = plt.subplots(figsize=(6.5, 5))
plt.rcParams['axes.facecolor'] = 'white'

dot_colors = cm.Greys(np.linspace(0.2, 0.7, len(datasets)))[::-1]
line_colors = cm.Reds(np.linspace(0.9, 0.2, len(datasets)))

datasets_sorted = sorted(datasets, key=lambda d: d["D"], reverse=True)

for i, dataset in enumerate(datasets_sorted):
    A, F, D = dataset["A"], dataset["F"], dataset["D"]
```

```
    Fmax, Fmin = params_fit[2*i], params_fit[2*i+1]
    A_fit = np.linspace(0, max(A), 200)
    F_fit = quenching_model(A_fit, Kd_fit, D, Fmax, Fmin)
    ax.plot(A, F, 'o', color=dot_colors[i], markeredgecolor='black', markersize=5)
    ax.plot(A_fit, F_fit, '-', color=line_colors[i], linewidth=2.2,
            label=f'[D]={D:.3f} µM')

ax.set_xlabel('[Protein B (Acceptor)] (µM)', fontsize=13)
ax.set_ylabel('Fluorescence (RFU)', fontsize=13)
ax.set_title(f'Global Donor Quenching Fit\nKd = {Kd_fit:.3g} µM,  R² = {R2:.3f}', fontsize=13, pad=10)
ax.spines[['top', 'right']].set_visible(False)
ax.grid(True, linestyle='--', color='lightgray', alpha=0.5)

# Legend styling
legend = ax.legend(
    frameon=False,
    loc='upper right',
    fontsize=11,
    handlelength=2.5,
    handletextpad=0.5,
    borderpad=0.3,
    labelspacing=0.4
)

plt.tight_layout()
plt.savefig("global_quenching_fit_raw.png", dpi=600, bbox_inches='tight', transparent=False)
plt.show()

print(f"Global fitted Kd = {Kd_fit:.4g} µM")
print(f"R² = {R2:.4f}")
```

### Sample acceptor emission fitting:

```
import numpy as np

import matplotlib.pyplot as plt

from scipy.optimize import curve_fit

import matplotlib.cm as cm

# === Your datasets ===

A0 =.12

X0= .4

datasets = [

  {

    "A": A0,

   "X": np.array([X0, X0/2, X0/4, X0/8, X0/16, X0/32, X0/64, 0

]),

    "EmFRET":
np.array([1703.74729,1679.570657,1142.737928,921.492572,498.4516879,301.1631506,105.7788147,0])

  },

  {

    "A": A0/2 ,

    "X": np.array([X0, X0/2, X0/4, X0/8, X0/16, X0/32, X0/64, 0

]),

    "EmFRET":
np.array([660.1645357,658.083211,624.3953035,619.0703754,425.145531,246.0378141,88.61781036,0])

  },

{

    "A": A0/4,

    "X": np.array([X0/2, X0/4, X0/8, X0/16, X0/32, X0/64, 0]),

    "EmFRET": np.array([181.8727424,200.2372575,209.9147173,193.6256831,161.4678365,68.9716586,0,])
```

```
  }
  ,
 {
    "A": A0/8,
    "X": np.array([  X0/4, X0/8, X0/16, X0/32, X0/64, 0]),
    "EmFRET": np.array([122.76655525,124.4985067,66.29626895,92.17357617,56.17889386,0])
  }
]

# === Global normalization ===
global_max = max(np.max(d["EmFRET"]) for d in datasets)
for d in datasets:
  d["EmFRET_norm"] = d["EmFRET"] / global_max
  d["EmFRET_max_raw"] = np.max(d["EmFRET"])

# === Binding model ===
def binding_model(X, Kd, A, EmFRET_max):
  Y = EmFRET_max - EmFRET_max * (
    2 * Kd / (X - A + Kd + np.sqrt((X - A - Kd)**2 + 4 * Kd * X))
  )
  return Y / global_max  # normalize globally

# === Global model ===
def global_model(dummy_X, Kd, Em_mult):
  model = []
  for dataset in datasets:
    X = dataset["X"]
    A = dataset["A"]
    Em_max = Em_mult * dataset["EmFRET_max_raw"]
    model.extend(binding_model(X, Kd, A, Em_max))
  return np.array(model)
```

```
# === Fit ===

initial_guess = [0.01, 1.2]

lower_bounds = [1e-9, 0.5]

upper_bounds = [1e3, 5.0]

Y_all = np.concatenate([d["EmFRET_norm"] for d in datasets])

X_dummy = np.arange(len(Y_all))

popt, _ = curve_fit(global_model, X_dummy, Y_all,

                p0=initial_guess, bounds=(lower_bounds, upper_bounds))

Kd_fit, Em_mult_fit = popt

# === Fit metrics ===

Y_fit_all = global_model(X_dummy, Kd_fit, Em_mult_fit)

SS_res = np.sum((Y_all - Y_fit_all)**2)

SS_tot = np.sum((Y_all - np.mean(Y_all))**2)

R2 = 1 - SS_res/SS_tot

RMSE = np.sqrt(np.mean((Y_all - Y_fit_all)**2))

# === Plot ===

plt.style.use('default')

fig, ax = plt.subplots(figsize=(6.5, 5))

# Color schemes

dot_colors = cm.Greys(np.linspace(0.15, 0.7, len(datasets)))[::-1]

line_colors = cm.Reds(np.linspace(1.0, 0.4, len(datasets)))  # red to pink

datasets_sorted = sorted(datasets, key=lambda d: d["A"], reverse=True)

# Plot data and fits

for i, dataset in enumerate(datasets_sorted):
```

```
    X, Y, A = dataset["X"], dataset["EmFRET_norm"], dataset["A"]
    Em_max = Em_mult_fit * dataset["EmFRET_max_raw"]
    X_fit = np.linspace(min(X), max(X), 200)
    Y_fit = binding_model(X_fit, Kd_fit, A, Em_max)

    ax.plot(X, Y, 'o', color=dot_colors[i], markeredgecolor='black', markersize=5)
    ax.plot(X_fit, Y_fit, '-', color=line_colors[i], linewidth=2.5,
        label=f'[A]={A:.3f} µM')

# === Styling ===
ax.set_xlabel('[Protein B] (µM)', fontsize=13)
ax.set_ylabel('FRET / FRET$_{max}$ (global)', fontsize=13)
ax.set_ylim(-0.05, 1.05)
ax.set_xlim(left=0)
ax.set_title(f'Purifed spytag003-mScarlet Kd={Kd_fit:.3g} µM, R²={R2:.3f}', fontsize=13, pad=12)
ax.spines[['top', 'right']].set_visible(False)
ax.grid(True, linestyle='--', color='lightgray', alpha=0.6)

# Legend — compact, in top-left corner with line color patches
legend = ax.legend(
    frameon=False,
    loc='upper left',
    fontsize=12,
    handlelength=2.8,
    handletextpad=0.6,
    borderpad=0.3,
    labelspacing=0.4
)

plt.tight_layout()
plt.savefig("global_fit_16ul.png", dpi=600, bbox_inches='tight', transparent=False)
```

```
plt.show()
```

```
plt.show()
```
